# STAG3 promotes exit from pluripotency through post-transcriptional mRNA regulation in the cytoplasm

**DOI:** 10.1101/2024.05.31.595485

**Authors:** Sam Weeks, Dubravka Pezic, Martin Dodel, Kunal Shah, Amandeep Bhamra, Stephen Henderson, Silvia Surinova, Tyson Sharp, Faraz Mardakheh, Suzana Hadjur

## Abstract

STAG proteins are key regulators of the cohesin complex and are often linked to alterations in cell identity and disease. Among the mammalian STAG paralogs, STAG3 has been less extensively studied beyond its known roles in meiosis. In this work, we demonstrate that STAG3 is expressed in mouse embryonic stem cells (mESCs) and primordial germ cell-like cells (PGCLCs), where it is required for cell fate decisions. Distinct from the other STAG proteins, STAG3 mediates its effects in the cytoplasm, facilitating the post-transcriptional regulation of gene expression. Furthermore, STAG3 localises to the centrosome independently of cohesin and interacts with proteins involved in mRNA localisation and stability. The knockdown of STAG3 in mESCs using siRNAs results in the destabilisation of the centrosome and the key P-body RNA-induced silencing complex (RISC) component TNRC6C, leading to the derepression of P-body localised mRNAs, such as DPPA3. Our results propose a model in which STAG3 collaborates with RNA-binding proteins (RBPs) and specific target mRNAs to control post-transcriptional gene expression and facilitate the transition from pluripotency in mESCs. Given that STAG3 is upregulated in various cancers, our results provide a novel perspective on how STAG proteins might contribute to cell identity and disease.

## INTRODUCTION

Cell identity is established through a complex collaboration of transcriptional and post-transcriptional mechanisms to fine-tune gene expression. Cells employ many post-transcriptional mechanisms to control gene expression, including the localisation of mRNAs to distinct cellular organelles or sub-compartments for the spatial and temporal control of protein synthesis (Das et al., 2021). This highly conserved process creates structural and functional asymmetries in cell identity such as those needed during embryonic patterning, cell migration, and synaptic plasticity (Chin et al., 2020).

The targeting of mRNAs to specific subcellular sites requires both *cis*-acting elements in the mRNA (often within the 3’UTR) and *trans*-acting RNA-binding proteins (RBPs). The resultant ribonucleoprotein (mRNP) complex mediates diverse aspects of RNA and protein metabolism including localised translation and motor protein mediated transport along the cytoskeleton (Martin et al., 2009). Aggregates of different mRNPs form condensates which act as hubs for post-transcriptional regulation. For example, stress granules and P-bodies sequester mRNA transcripts or destabilise mRNAs via the microRNA-induced silencing complex (RISC), thereby repressing translation of the mRNA cargo (Teixeira et al., 2005; Hubstenberger et al., 2017). The dynamic and heterogenous nature of these condensates enables rapid storage and release of mRNAs to fine-tune translation in a cell-type specific manner. In addition to these well-known cytoplasmic condensates, it is increasingly clear that the centrosome also acts as a site for localised translation (Sepulveda et al., 2018). Further to its canonical roles in microtubule nucleation and cell cycle control, mRNAs, RBPs, ribosomes and stable P-body proteins (Aizer et al., 2008; Moser et al., 2011) have all been detected at the centrosome, pointing to it as a hub of post-transcriptional gene regulation (Zein-Sabatto et al., 2021).

STAG proteins are known to regulate cohesin’s functions in genome organisation and gene expression within the nucleus (Pezic et al., 2017). STAG proteins are commonly mutated in cancer and developmental disease (Hill et al., 2016; Romero-Pérez et al., 2019; Jay et al., 2021), emphasising their importance in cell identity, although we do not have a full understanding of the mechanisms by which these proteins function. Recently, we have described non-canonical, cohesin-independent roles for STAG1 and STAG2 in RNA binding, interaction with RBPs and stabilisation of RNA:DNA hybrids in both mouse and human cells (Porter et al., 2023). Furthermore, STAG1 plays an important role in the structure and function of nucleolar condensates in mouse embryonic stem cells (mESCs), influencing cell fate plasticity (Pezic et al., 2023). Our work points to a range of novel activities that STAG proteins perform, often without the core cohesin ring, to influence cell identity beyond transcriptional regulation.

Here we show that STAG3, considered to be the ‘meiosis-specific’ STAG paralog (Pezzi et al., 2000; Prieto et al., 2001), is a key regulator of pluripotency exit in mESCs by supporting post-transcriptional regulation of gene expression in the cytoplasm. Proteomic analysis reveals that STAG3 interacts with an array of granule-associated RBPs and is localised to discrete cytoplasmic bodies including the centrosome. Loss of STAG3 impacts centrosome structure with no effect on cell cycle, and destabilises TNRC6C, a key component of the RISC complex. STAG3 is required for post-transcriptional control of *Dppa3,* thereby ensuring proper exit from pluripotency. Our results reveal a novel role for STAG3 in the post-transcriptional regulation of mRNAs that control cell fate.

## RESULTS

### *Stag3* is expressed in mESCs and is required for the exit from pluripotency

We have previously shown that of the three STAG paralogs, *Stag1* levels are dominant in 2i-grown (naïve) mouse embryonic stem cells (mESCs), and that *Stag1* and *Stag2* differ in their relative levels during *in vitro* epiblast-like cell (EpiLC) differentiation (Pezic et al., 2023). *Stag3* is considered a meiosis-specific paralog, even though its expression profile outside of terminally differentiated germ cells is poorly understood. In agreement with a recent study (Choi et al., 2022), we found that *Stag3* is expressed in mESCs, where its mRNA levels are 3.9-fold and 2.4-fold lower than *Stag1* and *Stag2*, respectively (Fig 1a, S1a). STAG3 protein was readily detected in mESCs by Immunoblotting (IB), with a greater abundance in serum-grown compared to naïve mESCs (Fig1b, S1b). Both *Stag3* mRNA and protein expression increased during EpiLC differentiation in a pattern similar to *Stag2,* a known pro-differentiation marker (Viny et al., 2019), consistent with a potential role for *Stag3* in the exit from pluripotency (Fig. 1b, S1c).

**Figure 1.**
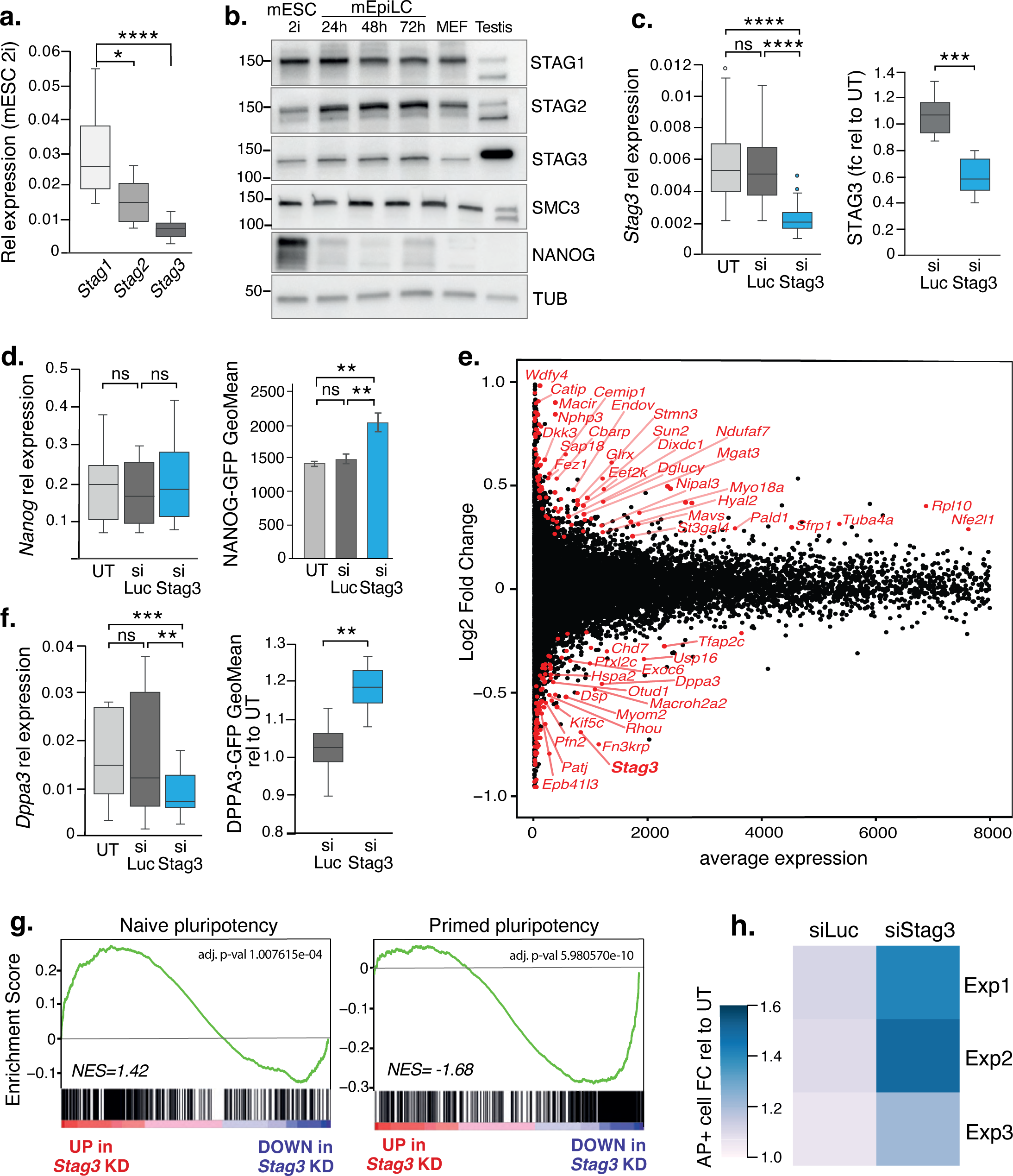
mESCs require *Stag3* to exit pluripotency. a) Relative mRNA expression of the three *Stag* paralogs by qRT-PCR in naïve (2i) mESCs. Data is from 15-20 independent biological replicates. The central line represents the median. Asterisks indicate a statistically significant difference as assessed using two-tailed t-test. *p < 0.05, **p < 0.005, ***p < 0.0005, ****p < 0.0001; ns, not significant. b) Whole-cell protein extracts (WCE) from mESCs, at several timepoints during EpiLC differentiation, in mouse embryonic fibroblast cells (MEF) and in Testis tissue were analysed by immunoblot (IB) for levels of STAG1, STAG2, STAG3, SMC3 and NANOG. alpha-TUBULIN (aTUB) serves as a loading control. c) Left, relative mRNA expression of *Stag3* by qRT-PCR in untreated (UT) mESCs or upon treatment with siLuc or siStag3. Data is from 20 independent biological replicates. Right, quantification of STAG3 protein levels in naive mESCs treated with siLuc and siStag3 assessed by ImageJ relative to the signal in UT cells. Data is from 5 independent biological replicates. Quantifications and statistical analysis as above. d) Left, relative mRNA expression of *Nanog* by qRT-PCR in UT, siLuc and siStag3 treated mESCs. Data is from 12 independent biological replicates. Right, quantification of NANOG protein levels in naive mESCs treated as above assessed by FACS analysis of NANOG-GFP GeoMean. Data is from 3 independent biological replicates. e) MA plot of RNA-seq obtained from 3 biological replicates of siStag3 and siLuc treated naive mESCs. Differential expression analysis was performed to plot data is as log2 fold change in KD conditions relative to siLuc controls. Labelled genes represent statistically significant (p<0.05) fold change based on students t-test. f) Left, relative expression of *Dppa3* mRNA by qRT-PCR in UT mESCs or upon treatment with siLuc or siStag3. Data is from 12 independent biological replicates. Right, quantification of DPPA3 protein levels in naive mESCs treated as above assessed by FACS analysis of DPPA3-GFP GeoMean relative to UT. Data is from 5 independent biological replicates. g) Enrichment score (ES) plots from gene set enrichment analysis (GSEA) using curated naive or primed pluripotency gene sets (see Methods). Negative and positive normalised enrichment scores (NES) point to the gene set being over-represented in the top-most down-or upregulated genes in *Stag3* KD mESCs, respectively. Vertical bars refer to individual genes in the gene set and their position reflects the contribution of each gene to the NES. h) Area occupied by AP colonies relative to total colony area in mESCs treated with siLuc and siStag3 from 3 independent biological replicates where n > 50 colonies/condition were counted.

To investigate the functional importance of *Stag3* in pluripotency, we established a *Stag3* knockdown (KD, ‘siStag3’) strategy using siRNAs (Methods). siRNA-mediated KD in naïve mESCs resulted in a mean reduction of *Stag3* mRNA by 62% compared to both untreated (UT) and cells treated with control siRNAs to Luciferase (siLuc) (Fig 1c). This was reflected in a mean reduction of STAG3 protein by 58% (Fig 1c, S1d). We observed no significant effect on the abundance of the other *Stag* paralogs, the cell cycle, or cell death (Fig S1e-h). qRT-PCR of the key pluripotency transcription factors *Nanog, Oct4, Sox2* and *Klf4* revealed no significant differences upon *Stag3* KD compared to controls (Fig 1d, S1i). Surprisingly, NANOG protein levels were increased by 44% upon *Stag3* KD in both NANOG-GFP mESCs as well as in WT mESCs (Fig 1d, S1j), revealing a difference in *Nanog* mRNA and protein regulation, and a possible role for STAG3 in pluripotency.

To measure the impact of *Stag3* KD on the global mESC transcriptome, we prepared RNA-sequencing libraries from three biological replicates of untreated, siLuc and siStag3 mESCs and used differential expression analysis to compute a log2 fold-change for each gene (Table S1). We found that most deregulated genes were lowly expressed in UT and siLuc mESCs (average normalised counts<500), and very few genes with average normalised counts >1000 were significantly affected by *Stag3* KD (Fig 1e). Overall, 54 genes were significantly up- and 40 genes were downregulated respectively by at least 1.5-fold compared to siLuc controls. In agreement with our previous results (Fig 1d, S1i), pluripotency transcription factors were not affected upon *Stag3* loss. One notable exception was *Dppa3 (Stella)*, a known regulator of both pluripotency and primordial germ cell (PGC) commitment (Sato et al., 2002), which was significantly reduced upon *Stag3* KD. Interestingly like NANOG, DPPA3 protein levels were increased (Fig 1f), revealing an unexpected disconnect between mRNA and protein levels of key cell fate regulators upon loss of *Stag3* in mESCs.

Using Gene Set Enrichment Analysis (GSEA) of custom gene lists (Pezic et al., 2023), we assessed the impact of *Stag3* KD on naïve and primed pluripotency signatures. In contrast to *Stag1,* which is required to maintain the naïve state, *Stag3* KD in mESCs revealed an enrichment for the naïve pluripotency-associated signature and a reduction of the primed pluripotency signature (Fig. 1g). This is in line with a role for STAG3 in the exit from pluripotency, similar to previous observations for STAG2. Supporting this, *Stag1* KD in naive cells is concomitant with a significant increase in *Stag3* expression (Fig S1k). Finally, serum-grown mESCs were plated in self-renewal conditions at clonal density and the proportion of undifferentiated cells upon *Stag3* KD was determined by measuring the area occupied by the colonies with high alkaline phosphatase activity (AP+). In siLuc controls, 52% of plated cells displayed high AP+ staining, which was not different from UT cells, and thus retained their naive state. Upon *Stag3* KD, the proportion of AP+ colonies increased by an average of 41% compared to siLuc controls (Fig. 1h, S1l), indicating that mESCs have an enhanced ability to self-renewal upon *Stag3* KD.

### A pro-pluripotency phenotype is maintained during Embryoid Body differentiation upon *Stag3* KD

STAG3 is expressed in terminally differentiated oocytes and sperm (Fig 1b) where it is required for germ cell meiosis (Garcia-Cruz et al., 2010; Winters et al., 2014; Hopkins et al., 2014; Llano et al., 2014; Liu et al., 2021). Germ cells are specified early in development from Primordial germ cells (PGCs) and this process can be modelled *in vitro* using defined culture conditions whereby competency for PGC-like cell (PGCLC) identity emerges by 48hr of EpiLC differentiation (Hayashi et al., 2011). Little is known about the role of STAG3 in PGCs. Several pieces of evidence suggested that STAG3 expression in mESCs could be important for PGC specification. For example, STAG3 protein levels increased post-EpiLC differentiation as cells approached PGCLC competency (Fig 1b, S1b, c); unbiased GSEA of our *Stag3* KD RNA-seq data identified a significant depletion of a Spermatogenesis gene signature (Fig 2a); and several genes associated with PGC specification were significantly downregulated in mESCs upon *Stag3* KD, including the key PGC regulators *Nanos3, Tfap2c* and *Dnd1* (Fig 2b).

**Figure 2.**
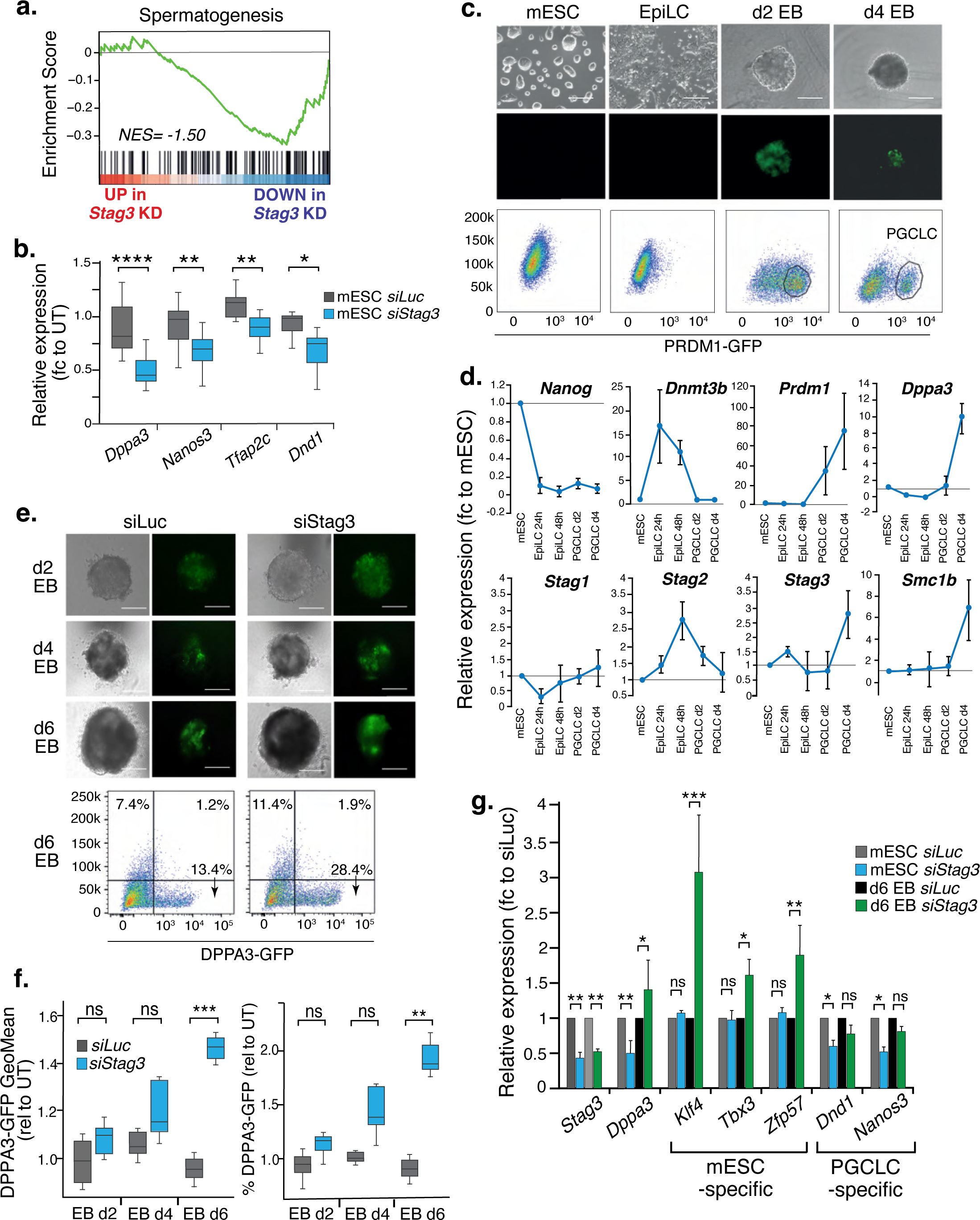
Loss of *Stag3* prevents commitment to PGCLC. a) GSEA of *Stag3* KD RNA-seq data reveals a negative normalised enrichment score of -1.50 (adj p= 0.044) for the Spermatogenesis signature (see Methods). b) Relative mRNA expression of PGC regulators in siLuc and siStag3 mESCs. Data is shown as fold-change from UT and is from 9 independent biological replicates. Quantifications and statistical analysis as before. c) Representative brightfield (top row), epi-fluorescent (middle row) and FACS profiles of PRDM1-GFP mESCs at select timepoints during *in vitro* differentiation into PGCLCs. Scale bar is 200um. d) Relative mRNA expression of select mESC (*Nanog),* EpiLC (*Dnmt3b*), PGCLC (*Prdm1, Dppa3*) and cohesin (*Stag1, 2, 3 and Smc1b*) genes from various timepoints during PGCLC differentiation *in vitro*. EB GFP+ cells were FACS sorted at the timepoints shown and collected for qRT-PCR. Gene expression is relative to levels in mESCs. Data is from 3 independent biological replicates. e) (Top) Representative brightfield and epi-fluorescent images of DPPA3-GFP mESCs treated with siLuc or siStag3 at three timepoints of embryoid body (EB) differentiation. (Bottom) Representative FACS profile of DPPA3-GFP d6 EB treated with siLuc or siStag3. Scale bar is 200μm. f) Left, quantification of DPPA3-GFP GeoMean or right, the % of DPPA3-GFP+ cells assessed by FACS (relative to UT mESCs) at different stages of PGCLC differentiation and upon siRNA treatment. Data is from 4 independent biological replicates. Quantifications and statistical analysis as before. g) Relative mRNA expression of select genes in mESCs or d6 EB GFP+ cells in siLuc and siStag3 conditions. Data is shown as fold-change from UT and is from 3 independent biological replicates. Quantifications and statistical analysis as above.

To study a potential role for STAG3 in PGC specification in mESCs, we established an *in vitro* PGCLC differentiation system using published protocols (Hayashi et al., 2011, and Methods). PGCLC specification was monitored using PRDM1-GFP (Ohinata et al., 2005) and DPPA3-GFP (Bleckwehl et al., 2021) mESCs. We tracked the emergence of GFP+PGCLCs using Fluorescence-activated cell analysis (FACS) and purified cells at various timepoints to measure gene expression changes within the GFP+ population (Fig 2c, d). As expected, within 24hr of EpiLC differentiation, *Nanog* levels decreased and *Dnmt3b* levels increased. Concomitant with PGC hypomethylation, embryoid body (EB)-containing PGCLCs had completely lost expression of *Dnmt3b* and significantly upregulated the known PGC markers, *Prdm1* and *Dppa3* (Fig 2d). As previously shown (Pezic et al., 2023), *Stag1* and *Stag2* have reciprocal expression profiles during early EpiLC differentiation (Fig 1b, 2d, S1c), with *Stag1* levels recovering somewhat in d4 PGCLCs. In contrast, both *Stag3 and Smc1b*, the germ-cell-specific component of the core cohesin complex (Revenkova et al., 2001), exhibited an increase in expression that was reminiscent of the germ-cell markers *Dppa3* and *Prdm1* (Fig 2d), suggesting a potential role for *Stag3* in early germ cell development.

To determine if *Stag3* is required for PGC specification, we treated both cell lines with *Stag3* and siLuc siRNAs and differentiated them towards PGCLCs as above. We established a KD protocol to ensure that *Stag3* levels were reduced as cells entered the EpiLC state, the timepoint that PGCLC commitment takes place *in vitro*. We collected samples from siLuc and siStag3 cells at multiple timepoints and used FACS analysis as above to monitor the impact on PGCLC development. Using the DPPA3-GFP mESCs, both the mean fluorescence of DPPA3-GFP and the frequency of GFP+ cells in the population were increased at every stage of EB differentiation after *Stag3* KD (Fig 2e, f). This was unexpected given the effect of *Stag3* KD on PGC marker expression (Fig 2b). Thus, we sought to validate this result by repeating the experiment in PRDM1-GFP cells. Unexpectedly, we found no significant difference in PGCLC numbers or PRDM1-GFP mean fluorescence upon *Stag3* KD using this cell line (Fig S2a-c).

As *Dppa3* is known to have roles in both pluripotency and PGCs, we hypothesised that the increased level of DPPA3-GFP in *Stag3* KD d6 EB may reflect sustained mESC-like identity (akin to Fig 1h), despite PGCLC culture conditions. To test this, we collected RNA from DPPA3-GFP mESCs and GFP+ d6 EB after treatment with siRNA to *Stag3* and performed qRT-PCR analysis using a panel of genes previously shown to be specific for naïve mESCs (*Klf4, Tbx3, Zfp57)* or PGCLCs (*Nanos3, Dnd1*) (Bleckwehl et al., 2021) (Fig 2g). Indeed, the GFP+ cells from d6 EB had significantly increased levels of mESC-specific mRNAs after *Stag3* KD compared to siLuc controls whilst the PGCLC-specific mRNAs were not significantly affected, reflecting a hyperESC-like state as opposed to PGCLC identity. Thus, despite a signalling environment that supports PGCLC commitment, *Stag3* KD cells maintained mRNA expression of mESC-specific genes and failed to fully commit to PGCLC identity.

### STAG3 is localised to the cytoplasm in mESCs

To begin to understand how STAG3 contributes to pluripotency, we extracted cytoplasmic, nuclear and chromatin protein fractions from mESC and EpiLCs and assessed STAG3 abundance in these fractions using immunoblotting. As before, we observed more STAG3 signal in EpiLCs compared to mESCs. Surprisingly, the majority of STAG3 protein was detected in the cytoplasmic fraction in both cell types and in both G1 and G2 phases of the cell cycle (Fig 3a, b). This was decidedly different from STAG1, STAG2 and RAD21 which were predominantly chromatin-associated in mESCs, as expected (Fig S3a). We did detect a weak signal of STAG3 on chromatin in mESC and EpiLCs, which was associated with a shift in size (Fig 3a, arrows in over-exposed IB). This matched the molecular weight of STAG3 detected in testis whole cell lysate (Fig 1b) and possibly reflects a bandshift due to phosphorylation of chromatin-bound STAG3 (Fukuda et al., 2012). Immunoprecipitation (IP) of chromatin-bound STAG3 enriched the SMC3 and RAD21 components of the cohesin ring and CTCF (Fig S3b), similar to a previous report (Choi et al., 2022) and in support of the idea of complex cohesin regulation in mESCs (Pezic et al., 2017; Pezic et al. 2023).

**Figure 3.**
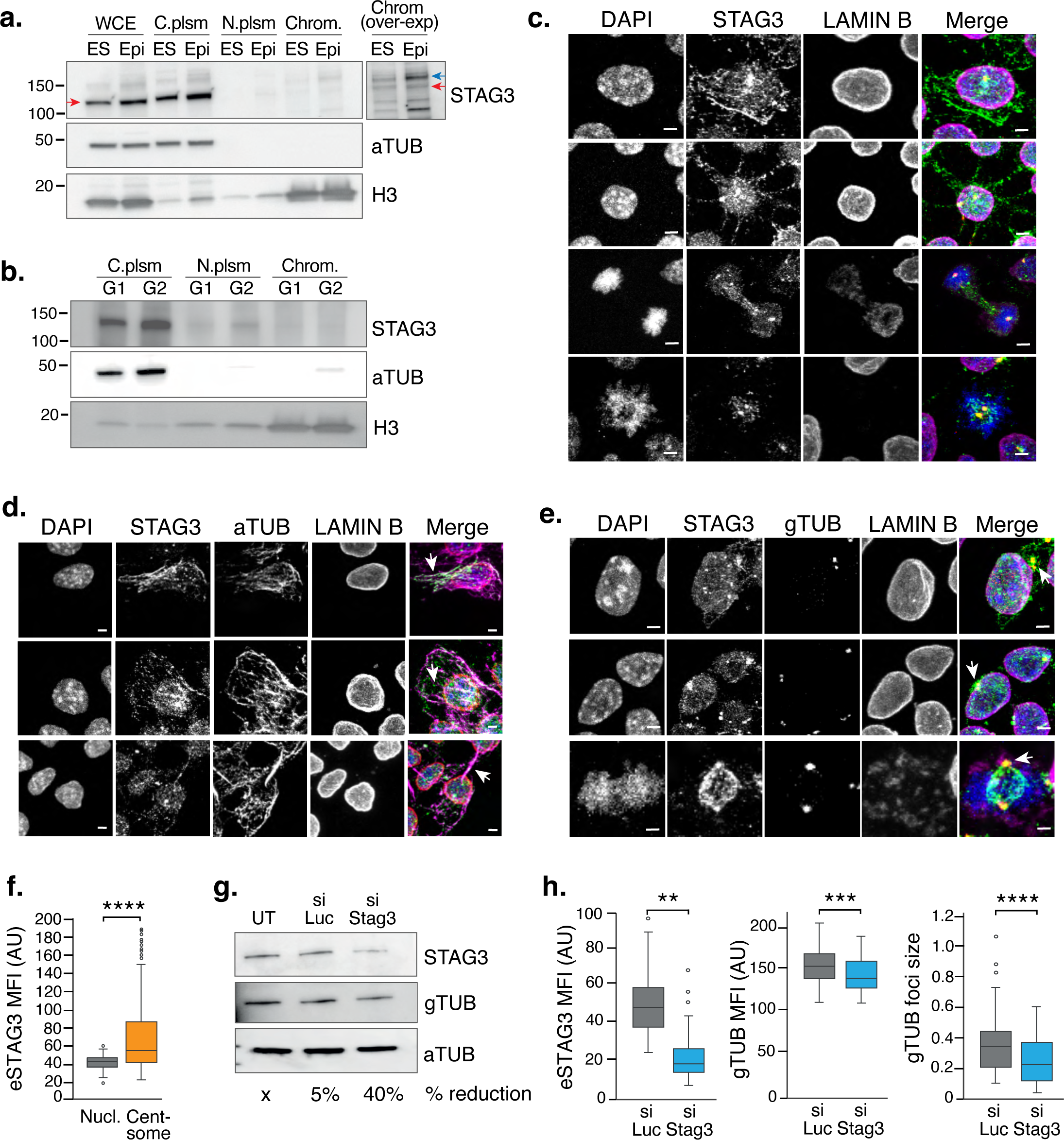
STAG3 is localised to the cytoplasm in mESCs. a) Immunoblot analysis of STAG3 levels in whole cell extract (WCE), cytoplasmic (C.plsm), nucleoplasmic (N.plsm) and chromatin (Chrom.) fractions in mESC (ES) and 48hr EpiLCs (Epi). Inset, over-exposed IB of the chromatin fraction to show the STAG3 bandshift (compare red and blue arrows). alpha-TUBULIN (aTUB) and H3 serve as fractionation and loading controls. b) Immunoblot analysis of STAG3 levels in fractionated lysates from FACS-sorted G1 or G2 mESC populations. aTUB and H3 serve as fractionation and loading controls as above. c) Representative confocal images of endogenous LAMIN B (demarcating the nuclear membrane) and STAG3 assessed by co-IF in mESCs and counterstained with DAPI. Shown are four independent cells with different STAG3 signal profiles in the cytoplasm. Scale bars, 3µm. d)-e) Representative confocal images of d) aTUB or e) gamma-(g-)TUBULIN (gTUB) with STAG3 and LAMIN B and counterstained with DAPI by co-IF in mESCs. Arrows indicate notable regions of overlap with STAG3 in the cytoplasm. *NB.* the STAG3 signal at the aTUB ‘bridge’ shown in the last cell from d). Scale bars, 3µm. f) ImageJ quantification of the MFI of endogenous (eSTAG3) in wildtype (WT) mESC in either the nucleus (light grey) or at gTUB foci (orange). Quantifications and statistical analysis are as shown previously. Data are from n > 100 independent cells/condition in 3 biological replicates. AU, arbitrary units. g) Immunoblot analysis of STAG3, gTUB and aTUB levels in WCE of UT mESCs or after treatment with siLuc or siStag3. Shown also is the % reduction in gTUB levels relative to the signal in UT cells, as assessed by quantification of the bands using ImageJ. h) ImageJ quantification of the MFI of left, eSTAG3; middle, gTUB and right, gTUB foci size in WT mESC treated with siLuc (grey) or siStag3 (blue). Quantifications and statistical analysis were done as above. Data are from n > 100 independent cells/condition in 2 biological replicates. AU, arbitrary units.

To validate these observations and to examine the pattern of STAG3 localisation in the cytoplasm, we performed Immunofluorescence (IF) of endogenous STAG3 (eSTAG3) protein in mESCs (Fig 3c, S3c, d, e). As expected from the fractionations, neither STAG1 nor STAG2 had any appreciable cytoplasmic signal in mESCs (Fig S3d). In contrast, STAG3 had a punctate cytoplasmic signal with three distinct patterns in mESCs; one large puncta often closely associated with the outer nuclear membrane and reminiscent of centrosomes; fibril-like extensions resembling the cytoskeleton; and, in cells undergoing mitosis, STAG3 was associated with the spindle (Fig 3c). Interestingly, the cytoplasmic STAG3 localisation appeared to be independent of RAD21 (Fig S3e), suggesting that STAG3 in the cytoplasm may be independent of the mitotic cohesin complex.

As the profile of STAG3 signal in mESCs was reminiscent of the cytoskeleton and centrosomes, we performed co-IF experiments with antibodies to alpha-(a-) and gamma-(g-)TUBULIN (TUB). We observed co-localisation of STAG3 with a-TUB fibrils throughout the cytoplasm, as well as at dense patches of a-TUB bridges between mESCs, a feature that is common in cells with high developmental potential and essential for germ cell maturation (Zenker et al., 2017; Hawdon et al., 2021) (Fig 3d). Similarly, STAG3 strongly co-localised with g-TUB foci in both non-dividing cells and at the spindle (Fig 3e, f). Fractionated mESCs did not reveal differences in the total levels of a-TUB protein in cytoplasmic extracts upon *Stag3* KD, however, we did observe a 40% reduction in total g-TUB protein levels (Fig 3g). As validation, we used CRISPR-Cas9 to knock-in a v5 tag into the 3’terminus of the endogenous *Stag3* locus to generate a heterozygous *Stag3*-v5 mESC clone (Fig S3f). STAG3-v5 was sensitive to siRNA-mediated *Stag3* KD, was localised to the cytoplasm and was enriched at g-TUB foci, confirming the phenotype in non-targeted, wildtype (WT) mESCs (Fig S3g, h). We assessed the impact of *Stag3* KD on centrosome structure by repeating the coIF experiment in siLuc and si*Stag3* conditions in both WT mESCs and our *Stag3*-v5 mESC clone. In both cell types, the g-TUB mean fluorescence signal and foci size were significantly reduced upon *Stag3* KD (MFI reduced by 8% in eSTAG3 and 6% in *Stag3*-v5, p<0.001; foci size reduced by 29% eSTAG3 and 21% in *Stag3*-v5, p<0.001) (Fig 3h, S3h), suggesting a role for STAG3 in centrosome stability in mESCs.

### STAG3 interactome in mESCs enriches for proteins associated with cytoplasmic condensates

We characterized the STAG3 protein–protein interaction (PPI) network in mESCs using Immunoprecipitation (IP) coupled with mass-spectrometry (IP-MS). Our repeated attempts at IP-MS of eSTAG3 with commercial antibodies proved unsuccessful (see Methods). Instead, we took advantage of our *Stag3-*v5 mESC clone and used v5-TRAP to characterize the STAG3-v5 PPI network in mESCs. Three biological replicates were prepared from *Stag3-*v5 and non-targeted (WT) mESCs and processed for IP. 337 immunoprecipitated (IP) proteins were identified by liquid chromatography tandem mass spectrometry (LC-MS/MS), with STAG3 peptides robustly detected in all *Stag3-*v5 mESC samples and never detected in WT mESCs, validating the specificity of the v5-TRAP. We compared the v5 IP to WT samples to generate a log2 fold-change value for each putative interactor. From this, we discovered 39 proteins that were STAG3-v5-specific (Fig 4a, blue dots) as well as interactors that were significantly changed by at least 2-fold (p-value ≤ 0.05) compared to WT controls (Fig 4a, red dots), yielding a total of 148 STAG3-v5 interactors (including STAG3) in mESCs (Table S2).

**Figure 4.**
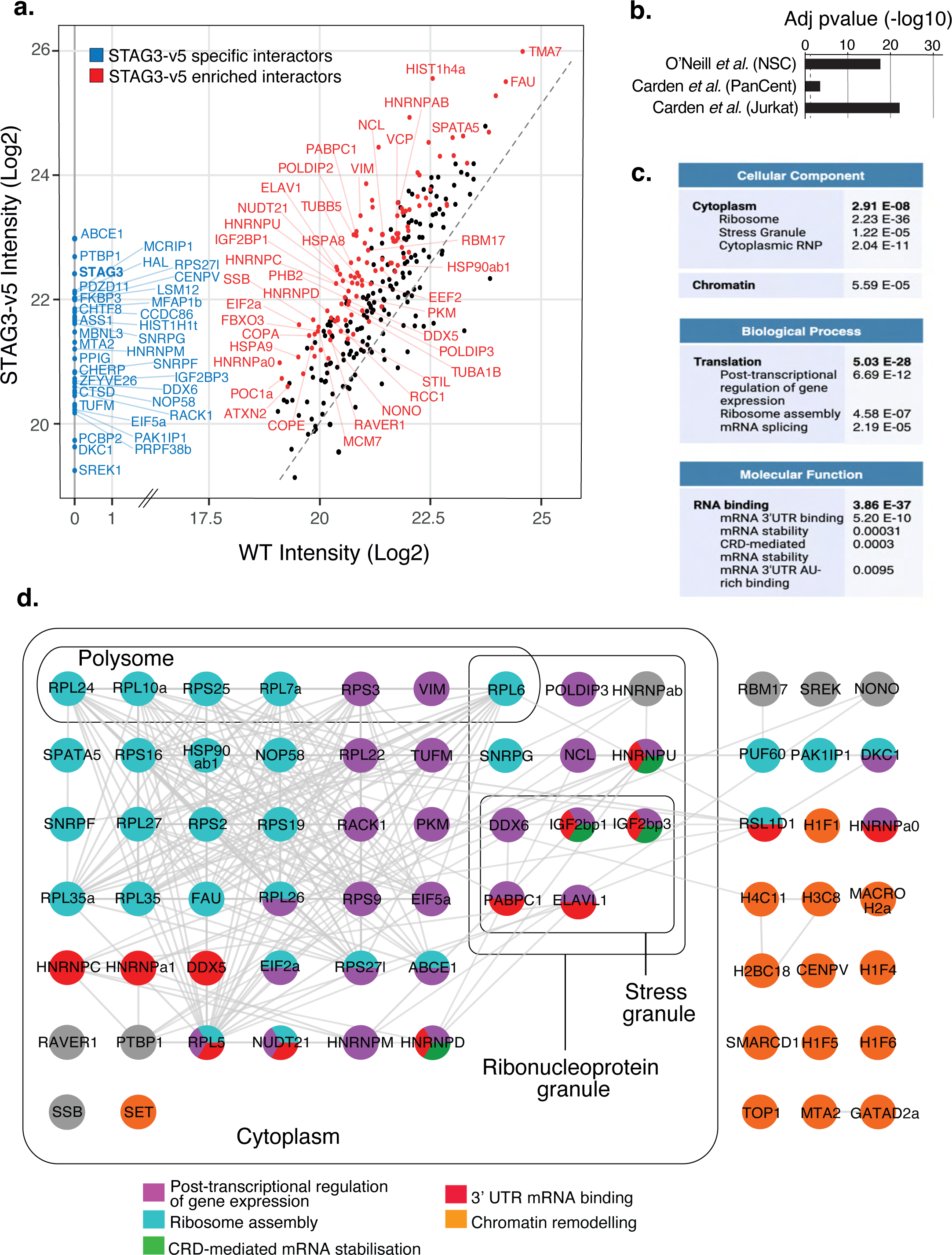
Characterisation of the STAG3 protein interaction network in mESCs. a) Scatter plot displaying the log2 protein group intensity across three biological replicates for the 337 proteins identified in STAG3-v5 IP-MS data produced from v5-TRAP in WT and *Stag3*-v5 mESCs. Coloured dots represent STAG3-v5 specific and enriched interactors (148 proteins in total). Blue dots/labels represent proteins which were uniquely detected in the v5 IPs, representing STAG3-v5-specific interactors. Red dots/labels represent enriched STAG3-v5 interactors with an abundance increase of at least 2-fold and p<0.05 in STAG3-v5 compared to WT*. NB.* For clarity we have not labelled all of the ribosome proteins here, please refer to Figure S4 and Table S2. b) -Log10 transformed adjusted p-value (FDR) for enrichment of centrosome interactome data from O’Neill *et al*. and Carden *et al*., with the STAG3-V5 interactome. ‘PanCent’ comprises centrosome-interacting proteins common to at least two of the four human cancer cell lines tested in Carden *et al*. c) Most enriched GO terms and their corresponding FDR values, arranged by category, for the 147 STAG3-v5 interactors (STAG3 was not included as an interactor). For full GO term enrichment, see Table S2. d) Simplified STAG3-v5 interaction network of protein–protein interactions identified in mESCs using STRING. Node colours describe the major enriched categories with squares denoting RBPs. Proteins are grouped according to *Cellular Compartment* enriched terms. All proteins present in both the 147 STAG3-v5 PPI and each enriched GO term are indicated here.

We used STRING to analyse associations in the STAG3-v5 interactome and to identify Gene Ontology (GO) based enrichment of functional terms, relative to the whole murine proteome. In agreement with our results that STAG3 is primarily a cytoplasmic protein in mESCs (Fig 3), we identified ‘Cytoplasm’ as an enriched term within *Cellular Compartment* GO in the STAG3-v5 PPI network. Further to our observations of STAG3 at the centrosome (Fig 3), ‘Ribonucleoprotein granules’, specifically including the ribosome, stress granules and paraspeckles were also *Cellular Compartment* enriched GO terms, suggesting STAG3 is localised to additional cytoplasmic condensates (Fig 4c, Table S2). Also, in support of our IF results, ‘Spindle organisation’ was a specifically enriched term and the key centriole assembly proteins POC1A and STIL were present in the STAG3 interactome. Several cytoskeleton interactors were enriched within the STAG3-v5 PPI network including VIM, TUBB5, TUBA1b, although we anticipate that our stringent IP conditions (Methods) may have precluded detection of more cytoskeleton interactors.

### STAG3 interacts with RNA regulators at the centrosome

Given the co-localisation of STAG3 with gTUB foci by IF (Fig 3e, f, S3h), we were surprised to not detect enrichment of traditional centrosome GO terms such as Microtubule organising centre (GO:0005815) or Centrosome (GO:0005813) (Bauer et al., 2016; Carden et al., 2023). To investigate this further, we looked for overlap of our STAG3-v5 PPI network with a recent human neural stem cell (NSC) centrosome proteome (O’Neill et al., 2022) and assessed the significance using a hypergeometric distribution (Methods). There was significant enrichment of STAG3-v5 interactors in the NSC centrosome dataset (24 proteins, FDR 3.7e-20), 10 of which were specific to the STAG3-v5 IP (Fig 4b, Table S2). Notably, only a few of the overlapping proteins were canonical centrosomal proteins (Bauer et al., 2016). Instead, the majority of proteins had functions in mRNA binding and stability (Table S2).

We validated this using a second recently published centrosome proteome derived from several human cell lines (Carden et al., 2023). As before, we found significant enrichment of centrosome-associated proteins from all the published cell lines in the STAG3-v5 PPI network, including the ‘pan-centrosome’ proteome (FDR 0.0019) (Fig 4b). Interestingly, the Jurkat centrosome proteome was the most significantly enriched (FDR 1.8 e-25) in the STAG3-v5 network (Fig 4b) and the only cell line with ‘Ribonucleoprotein complex’ enriched as a GO term (Carden et al., 2023), similar to the enrichment of mRNA regulation terms from the O’Neill proteome (Table S2). These observations are reminiscent of recently described centrosome-based roles in post-transcriptional gene regulation and localised translation (Zein-Sabatto et al., 2021). Together, this suggested that the localisation of STAG3 at the centrosome in mESCs may not be for centrosome biology *per se*, but rather for the purposes of post-transcriptional mRNA regulation at this location.

To determine if mRNA regulation was a general feature of the STAG3-v5 interactome, we returned to STRING to analyse enrichment of GO biological processes and molecular functions within the whole network. ‘Translation’ was significantly enriched as a *Biological Process* and included proteins involved in ‘post-transcriptional regulation of gene expression’, ‘Ribosome assembly’ and ‘mRNA splicing’ (Fig 4c, d, Table S2). ‘RNA binding’ was the most significantly enriched *Molecular Function* term and included proteins involved in ‘mRNA 3’UTR binding’, ‘mRNA stabilization’ and ‘CRD-mediated mRNA stabilization’ (Fig 4c, d). Finally, *SMART domain* database revealed a significant enrichment for RNA binding motifs (Methods). Taken together, our results pointed to a role for STAG3 in mRNA regulation and stability in the cytoplasm to control translation. Further, they suggested that STAG3 may be mediating these effects at cytoplasmic condensates such as centrosomes.

### STAG3 regulates *Dppa3* levels post-transcriptionally

A role for STAG3 in post-transcriptional regulation of gene expression could explain the disconnect we observed previously between mRNA and protein levels - whereby mESCs treated with siStag3 exhibited a reduction in *Nanog* and *Dppa3* mRNA but an increase in their protein levels compared to controls (Fig 1d, e, f, S1j). Indeed, *Dppa3* mRNA is known to be regulated post-transcriptionally by DDX6-mediated sequestration in cytoplasmic P-bodies (DiStefano et al., 2019). Loss of DDX6 in ESCs leads to unusually high levels of DPPA3 and consequently a hyper-ESC state, despite active differentiation signalling (DiStefano et al., 2019). This phenotype is remarkably similar to the enhanced pluripotent state (Fig 1g, h) and the maintenance of naïve pluripotency despite growth in PGCLC conditions (Fig 2g) that we observed upon *Stag3* KD. Furthermore, DDX6 and IGF2BP1/2 are known P-body proteins and are STAG3-v5 specific interactors (Fig 4a). Thus, we reasoned that STAG3 may negatively regulate *Dppa3* translation in mESCs and thereby control exit from pluripotency, akin to its interacting partner DDX6.

To test this, we treated mESCs with siStag3 or siLuc followed by cyclohexamide (CHX) to inhibit translation and then assessed the impact on *Dppa3* mRNA levels. *Stag3* was significantly reduced throughout the experiment (60% reduced, p<0.005 at each timepoint) and, as observed previously, siStag3 led to reduced *Dppa3* mRNA (51% reduced, p<0.005) and increased DPPA3 protein levels (Fig 1f, Fig 2b). Treatment with CHX rescued *Dppa3* mRNA in a time dependent manner, reaching levels similar to siLuc treated cells by 8hrs without evidence of cell death (Fig 5a). The rescue of *Dppa3* mRNA upon CHX treatment in *Stag3* KD reveals an unexpected role for STAG3 in negative regulation of *Dppa3* translation. This was not reflected in a change to global translation in mESCs treated with siStag3 (Fig. S5a), suggesting that STAG3 likely supports post-transcriptional regulation of specific mRNAs. Indeed, re-analysis of the genes deregulated upon *Stag3* KD (Fig 1e) revealed significant overlap with MSigDB gene sets representing Cytoskeleton Organisation (FDR 2.66E-02), Actin Filament Process (FDR 8.61E-05) and Regulation of Intracellular transport (FDR 6.51E-05) (Table S1), raising the possibility that STAG3 may play a role in post-transcriptional regulation of mRNAs required for the cytoskeleton and cytoplasmic trafficking.

**Figure 5.**
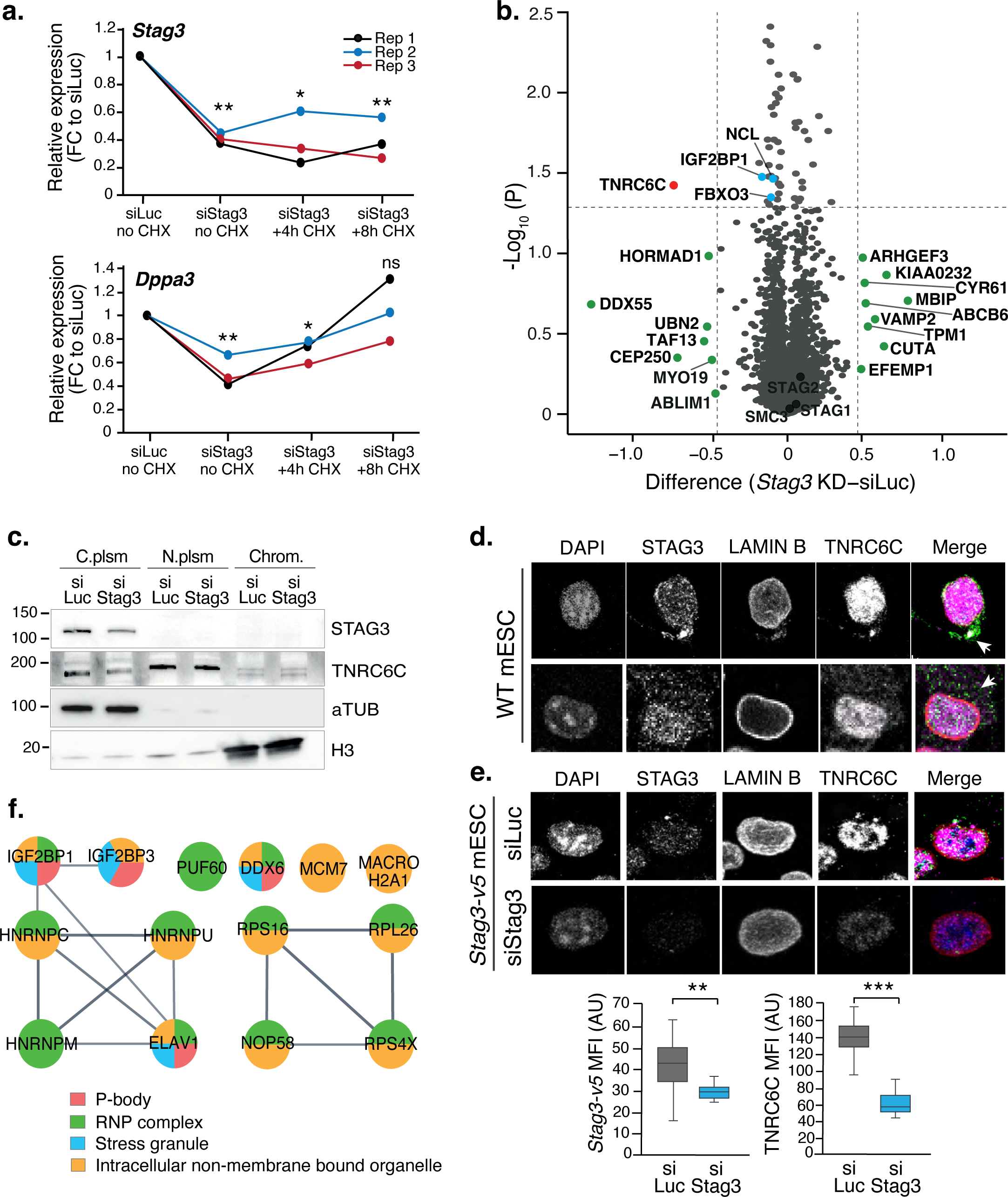
STAG3 mediates post-transcriptional regulation of *Dppa3* in mESCs and destabilises TNRC6C. a) Relative mRNA expression of *Stag3* (top) and *Dppa3* (bottom) in siLuc and siStag3 mESCs treated with cyclohexamide (CHX) for 4 or 8 hours to inhibit translation from three independent replicates. Significance refers to the average expression at each timepoint using t-test as before. b) Scatter plot from the TMT analysis displaying the statistical significance (-log10 p-value) versus the protein abundance difference from siLuc and siStag3 mESCs. Data was derived from three biological replicates. Vertical dashed lines represent changes of 1.5-fold (green dots). Horizontal dashed line represents a p-value of 0.05. Blue dots, STAG3-V5 interactors from Fig 4a. c) Immunoblot analysis of STAG3 and TNRC6C levels in cytoplasmic (C.plsm), nucleoplasmic (N.plsm) and chromatin (Chrom.) fractions of mESCs treated with siLuc and siStag3. aTUB and H3 serve as fractionation and loading controls. d) Representative confocal images of coIF of endogenous STAG3, LAMIN B and TNRC6C in WT mESCs treated with siLuc and siStag3. Arrows indicate notable regions of overlap. e) (Above) Representative confocal images of coIF of STAG3-v5, LAMIN B and TNRC6C under siLuc and siStag3 conditions. (Below) ImageJ quantification of the MFI of left, STAG3-v5 and right, TNRC6C in siRNA-treated mESCs. Quantifications and statistical analysis were done as above. Data are from n > 100 independent cells/condition in 2 biological replicates. AU, arbitrary units. f) STRING analysis of the STAG3-V5 interactors enriched within the P-body interactome data from Hubstenberger *et al*. -log10 transformed adjusted p-values (FDR) are in the text. Node colours describe the major enriched categories of the enriched proteins.

### Loss of STAG3 in mESCs destablises proteins associated with the cytoskeleton, centrosome and RNA metabolism

To study how STAG3 might regulate translation and identify its protein targets, we used Tandem Mass Tag (TMT) coupled with Mass spectrometry to unbiasedly measure the changes to the proteome of mESCs upon *Stag3* KD. TMT are chemical tags that allow samples to be multiplexed for relative quantitative proteomics analysis. We prepared three biological replicates of mESCs either untreated (UT) or treated with siStag3 and siLuc, and processed all 9 samples together with the TMT technique for differential proteome expression analysis. The experiment identified 5662 unique proteins in mESCs and we used a differential analysis to identify proteins whose abundance was sensitive to *Stag3* KD (Fig 5b, Methods, Table S3). No DPPA3 peptides were identified in our experiment, thus precluding us from assessing its expression level change via TMT, however NANOG revealed a small increase upon *Stag3* KD, reminiscent of our previous results (Fig 1d). As before, we found no effect on the abundance of any of the cohesin complex members, nor on the other STAG paralogs upon *Stag3* KD.

The majority of the TMT-detected proteins had small fold changes in abundance, with proteins both up and down regulated upon *Stag3* KD in mESCs (Fig 5b). 18 proteins showed a 1.5-fold change or greater and, although most of these destabilized proteins were not significantly changed, they were associated with biological functions that were in line with our previous IF and RNA-seq results. For example, HORMAD1, a known meiotic interactor of STAG3, is a component of the synaptonemal complex required for meiotic cross-overs and cohesion; DDX55 is an RNA helicase involved in translation initiation; CEP250 is a component of the centrosome and ABLIM1 mediates interactions between actin and cytoplasmic targets. Among these were genes whose mRNA levels were reduced (RNA-Seq), and protein levels were increased (TMT) upon *Stag3* KD, reminiscent of the post-transcriptional regulation of DPPA3 previously observed (Fig 5a). For example, TPM1 is associated with the cytoskeleton and VAMP2 plays a role in mRNA transport in vesicles. We validated that siStag3-repressed mRNA levels recovered upon inhibition of translation (as for DPPA3), for three proteins with roles in cytoskeleton and RNA regulation (Fig S5b). Together, these results point to a role for STAG3 in post-transcriptional regulation of select mRNAs.

### TNRC6C, a key regulator of P-body granules, is sensitive to STAG3 levels

The TMT assay indicated a significant reduction in the abundance of Trinucleotide repeat-containing gene 6C (TNRC6C) in mESCs treated with siStag3, compared to the control siLuc (a 73% decrease, p=0.038). TNRC6C is essential for post-transcriptional gene silencing via the RNA interference (RNAi) and microRNA pathways and, along with other components of the RISC complex, is involved in forming cytoplasmic granules with mRNAs, known as P-bodies. Additionally, a smaller yet significant reduction in insulin-like growth factor 2 mRNA-binding protein 1 (IGF2BP1) was observed following *Stag3* KD (15% reduction, p=0.033) (Fig 5b, blue dot, Table S3). IGF2BP1, which interacts with STAG3-V5 (Fig 4a), is also a component of P-bodies. Other RNA binding proteins similarly showed modest but significant protein reduction, such as STRAP (11% decrease, p=0.0038), EZH2 (10% decrease, p=0.012) and MAGOH (9% decrease, p=0.019). Indeed, GO analysis of significantly reduced TMT hits (p<0.05, 33 proteins) revealed an enrichment of *Biological Process* ‘Regulation of nuclear-transcribed mRNA catabolic process, deadenylation-dependent decay’ (FDR 0.014) and *Molecular Function* ‘RNA Binding’ (FDR 0.029) terms (Table S3), raising the possibility STAG3 stabilises RNPs involved in mRNA regulation. The decrease in TNRC6C levels was confirmed using two independent techniques. First, immunoblot analysis of mESCs treated with either siStag3 or siLuc followed by cellular fractionation, demonstrated a 47% reduction in TNRC6C in the cytoplasm (Fig 5c). Second, co-immunofluorescence (coIF) studies showed co-localisation of TNRC6C with STAG3 in both the nucleoplasm and cytoplasm of mESCs, where it was sensitive to siStag3 (Fig 5d, e, S5c, d). These observations are consistent with the findings from the TMT and IB assays.

Interestingly, cytoplasmic TNRC6C signal in both WT and *Stag3*-v5 mESCs appeared as both dispersed puncta and a distinct focus at the centrosome, where it colocalised with STAG3 (Fig 5d, S5c). Both TNRC6C signal profiles were sensitive to siStag3, indicating that the effect on TNRC6C is at least in part due to the destabilisation of centrosomes upon *Stag3* KD. Indeed, it is known that while most cytoplasmic P-bodies are mobile and can transit rapidly along microtubule tracks to and from the centrosome, stationary P-bodies reside at the centrosome in interphase cells (Aizer et al., 2008; Moser et al., 2011), where they may act as local translation hubs (Liu et al., 2023). This raised the question of whether STAG3 is a component of P-bodies and contributes to mRNA-mediated stabilisation in this context. In support of this, STAG3 interacted with DDX6, ELAVL1 and IGF2BP1/3 (Fig 4a), all of which play a role in post-transcriptional regulation at TNRC6-containing granules and at centrosomes. We thus analysed the overlap of P-body-associated proteins (Hubstenberger et al., 2017) with our STAG3-v5 PPI network and assessed significance as before (Methods). The P-body proteome was highly enriched in the STAG3-v5 interactome, identifying 15 overlapping proteins (hypergeometric distribution = 8.9e-21, FDR = 1.4e-19, Log10 fold enrichment = 1.65) (Fig 5f). Together our results suggest that STAG3 may repress the translation of specific mRNAs in P-bodies, either those transported along the cytoskeleton or stationary ones at the centrosomes.

## DISCUSSION

Here we report that the cohesin regulator STAG3 is not restricted to regulating cohesion in meiosis but is also expressed in pluripotent cells where it contributes to exit from the naïve pluripotent state. Unexpectedly, STAG3 mediates its effects in the cytoplasm where it colocalises with and destabilises the centrosome and cytoskeleton and interacts with a variety of RBPs required for post-transcriptional regulation of gene expression. We discover a specific role for STAG3 in regulation of *Dppa3* mRNA, which is required for cells to exit pluripotency. Our observations expand the role for STAG3 out of the nucleus and support the growing body of evidence that STAG proteins perform a variety of functions that extend well beyond the fine-tuning of cohesin on chromatin.

Like STAG2, STAG3 acts as a pro-differentiation marker in mESCs. Reduced levels of *Stag3* led to increased levels of DPPA3 and a ‘hyper-ESC’ state that was resistant to differentiation stimuli, as observed here and in (DiStefano et al., 2016). This epigenetic regulator prevents *de novo* DNA methylation whilst protecting the methylation status of imprinted genes in mESCs (Nakamura et al., 2012). Thus, siStag3-mediated DPPA3 up-regulation in mESCs may prevent the emergence of a hypermethylated state, which is required for primed EpiLCs and subsequent PGCLC commitment. It is noteworthy that *Stag3* expression also increased in EB, in line with other PGC regulators such as *Prdm1*. Thus, it remains to be seen if STAG3 plays different roles at distinct stages of PGC development.

The increased levels of DPPA3 and NANOG protein we observed upon *Stag3* KD did not reflect a global change in translation in mESCs. Such *selective* post-transcriptional regulation is in line with RBP-specificity for mRNA subsets that contain recognition sequences, such as AU-rich elements (AREs), the motif which is present in both *Nanog* and *Dppa3* 3’UTRs (Otsuka et al., 2019). Of the STAG3 interactors, several RBPs (DDX6, ELAVL1, HNRNPD) are known to bind ARE targets (Loflin et al., 1999; DiStefano et al., 2019; Bakheet et al., 2018), and cross-linking immunoprecipitation followed by RNA sequencing (CLIP-Seq) reveals that DDX6 directly binds *Nanog* mRNA (DiStefano et al., 2019). In this context, the centrosome also acts as a post-transcriptional regulator of gene expression (Woodruff et al., 2017) whereby specific 3’UTR ‘zip code’ sequences sequester select mRNAs to the centrosome for localised translation (Zein-Sabatto et al., 2021). Given that interphase centrosome structure becomes destabilised upon loss of *Stag3*, it is tempting to speculate that this effect is due to changes in specific mRNA localisation or translation at the centrosome mediated by STAG3 and its interacting RBPs.

We note that STAG3s influence on cell identity, such as in pluripotency and cancer, goes well beyond transcription factor function and chromatin regulation. Indeed, mRNAs with functions in cytoskeleton, vesicle trafficking and RNA regulation were rescued by CHX in *Stag3* KD conditions, raising the possibility that STAG3 contributes to post-transcriptional regulation of specific mRNAs involved in these cellular activities. Increased cell migration and metastasis are linked to STAG3 de-repression in cancer (Sasaki et al., 2021), raising the question of whether post-transcriptional mechanisms are deregulated in STAG3-associated cancers. Additional experiments are required to determine the full spectrum of STAG3 mRNA targets. Moreover, the specific mechanism(s) by which STAG3 represses mRNA translation are yet to be determined. For example, STAG3 could function indirectly by supporting the cytoskeleton framework needed for transport of mRNA or the cytoplasmic bodies themselves. Alternatively, STAG3 may directly bind mRNAs or their RBPs and facilitate condensate structure needed to repress translation. Indeed, the STAG3-v5 PPI included components of the RISC complex (IGF2BP2, DDX6 and LSM12), identified *CRD-mediated mRNA stabilization* (Fig 4c) and *P-bodies* (Fig 5f) as enriched categories and TNRC6C was itself significantly destabilised upon *Stag3* KD. Altogether this supports a hypothesis that STAG3 contributes to the RISC-mediated post-transcriptional program.

Finally, both STAG1 and STAG2 are known to interact with RNAs (Porter et al., 2023) and given the high degree of conservation between the paralogs, it is very likely that STAG3 also binds RNA directly. This raises the interesting possibility that cells have evolved multiple STAG paralogs which occupy distinct cellular locations in order to coordinate mRNA regulation throughout the mRNA life cycle. In such a model, nuclear STAG1/2 could bind nascent RNA and recruit RBPs for functions such as RNA stabilisation, splicing, and export. Upon translocation to the cytoplasm, STAG3 may then take over, either binding RNA directly or via its interacting RBPs, to facilitate RNA localisation and/or trafficking and post-transcriptional regulation of its target. Such a ‘tag-team’ approach would point to an exquisite evolutionary divergence in an ancestral STAG protein to enable precise regulation of specific mRNAs in different cell compartments to fine-tune cell identity.

## ACKNOWLEDGMENTS

This work was supported by the Wellcome Trust through a Senior Research Fellowship awarded to S.H. (106985/Z/15/Z); a BBSRC I-CASE studentship (BB/R506199/1, in partnership with B. Taylor at Astra Zeneca) and a LiDo DCD Fellowship to S.W. Work at the Cancer Institute Translational Technology Platforms was supported by the CRUK City of London Centre Award (C7893/A26233) and (CTRQQR-2021/100004). We thank A. Rada-Iglesias and T. Bleckwehl for DPPA3-GFP ESCs and PGCLC differentiation advice, A. Surani for the PRDM1-GFP ESCs and support with the PGCLC differentiation and S. Godinho for advice with cytoskeleton immunofluorescence. Thank you to Y. Guo and J. Manji in the Cancer Institute CRUK Center FACS and Imaging core facilities for their invaluable assistance. We are grateful to the members of the Hadjur lab for critical discussions throughout the project.

## AUTHOR CONTRIBUTIONS

S.W. and S.H. conceived the project. S.W. designed and performed all the experiments and most of the analyses, with assistance early on in the project from D.P. RNA-seq libraries were prepared by the CRUK Centre Genomics Facility and sequenced at UCL Genomics. M.D. and F.M. performed the TMT experiment with samples provided by S.W. and F.M. did the statistical analysis. STAG3-v5 mass spectrometry and analysis were done by A.B, with advice from S.S. K.S performed human STAG3 protein analysis and reporter assays, and T.S. offered advice. S.Henderson performed the initial RNA-seq analysis. S.W. and S.H. formatted all figures and wrote the manuscript.

### Accession Numbers

RNA-sequencing data generated in this study was submitted to GEO with the Accession GSE267691. The STAG3-v5 mass spectrometry proteomics data were deposited to the ProteomeXchange Consortium via the PRIDE partner repository with the dataset identifier PXD046456.

## METHODS

### Embryonic stem cell culture and EpiLC differentiation

Wildtype (WT) E14, Dppa3-T2A-GFP, Prdm1-GFP and Stag3-v5 mouse embryonic stem cells (mESCs) were cultured in serum (FCS/LIF) or naïve (2i/LIF) conditions as indicated. Serum-cultured cells were grown on 0.1% gelatin-coated plates in Serum ES media (GMEM, 10% FCS (Sigma), 2mM Glutamax, 1X NEAA, 1mM Sodium Pyruvate, 0.1mM ßMercaptoethanol (BMe)) and freshly added mouse recombinant LIF (1000U/ml; Stemcell technologies 78056). 2i-cultured mESCswere grown on plates coated with Fibronectin (17.6μl/1ml PBS) (1mg/ml; Millipore FC010), in ES 2i media (DMEM:F12/Neurobasal 1:1, 1X KnockOut Serum Replacement, 1X N2 supplement, 1X B27 supplement, 2mM Glutamax, 1X Pen-Strep, 10mM HEPES, 1µM PD0325901 (Stemcell technologies 72182), 3µM CHIR99021 (Stemcell technologies 72052), 0.1mM BMe) and freshly added LIF as above. Epiblast-like cell (EpiLC) induction was achieved by seeding mES (2i/LIF-grown) cells in EpiLC media (DMEM:F12/Neurobasal 1:1, 1X KnockOut serum replacement, 1X N2 supplement, 1X B27 supplement, 2mM Glutamax, 1X Pen-Strep, 10mM HEPES) freshly supplemented with Activin A (20ng/ml; Peprotech 120-14P) and bFGF (12ng/ml; Peprotech 100-18B) growth factors. Cells were harvested at 24h, 48h and 72h post-EpiLC induction.

Embryoid Bodies (EB) were grown in a suspension culture in U-bottomed 96-well plates (ASMBio), which were pre-treated with 0.1% Lipidure (ASMBio, AMS.52000011GB1G) in 100% ethanol (30μl/well) and left in a sterile hood over night to air-dry. At 48h post-EpiLC induction, cells were seeded at a density of 4x10^3^ cells/well in 50μl Primordial germ cell-like cell (PGCLC) media (GMEM 15% KnockOut serum replacement, 1mM Sodium Pyruvate, 1X NEAA, 2mM Glutamax, 1X PenStrep, 50mM BMe) supplemented with human recombinant BMP4 (500ng/ml; Proteintech HZ-1045) and mouse recombinant LIF (1000U/ml). EBs were collected at d2, d4, and d6 time-points. To cellularise, 20 pooled EBs were incubated in 1ml accutase for 12min in a 15ml falcon tube with gentle shaking before being quenched with wash buffer (GMEM, 15% KnockOut serum replacement).

### siRNA-mediated knockdown and qRT-PCR analysis

siRNAs were purchased from Horizon Discovery (previously Dharmacon), including the siLuc control. siRNA knockdowns (KDs) were performed for 48hr in 6-well plates where 200,000 cells were seeded and 50pmol siRNA was transfected using 7.5µl RNAiMax Lipofectamine in 300µl Optimem at the time of seeding. The next day, media was changed, and the cells were transfected a second time with an additional 50pmol siRNA. siStag3 ‘SmartPool’ (SP) was purchased from Dharmacon and consisted of an equal mix of 4 siRNAs targeting Stag3 exons 4, 6, 8 and 26 (L-043704-01-0050). For qRT-PCR analyses, total RNA was isolated using Monarch RNA prep kit (NEB). Reverse transcription was performed on 0.5µg DNase-treated total RNA using Lunascript RT (NEB) in 20µl reactions. qRT-PCR was performed using 2x SensiFAST SYBR No-ROX kit (Bioline) in 20µl reactions using 1µl of RT reaction as input and 0.4µM each primer.

### Protein Lysates, Fractionations and Immunoblotting

Whole cell extracts (WCE) were collected by lysis in RIPA buffer (150mM NaCl, 1% NP-40 detergent, 0.5% Sodium Deoxycholate, 0.1% SDS, 25mM Tris-HCl pH 7.4, 1mM DTT) (1x10^6^/100µl buffer) and sonicated at 4°C for x5 30 second cycles using Diagenode Bioruptor. Insoluble material was pelleted and the supernatant lysate was quantified using BSA Assay (Thermo Scientific). For cellular fractionations, a cellular ratio of 5x10^6^ cells/100µl buffer was maintained throughout the protocol. Cells were re-suspended in Cell Membrane Lysis Buffer (0.1% Triton X-100, 10mM HEPES pH 7.9, 10mM KCl, 1.5mM MgCl2, 0.34M sucrose, 10% glycerol, 1mM DTT), incubated on ice for 5min and centrifuged for 5min at 3700rpm to collect the cytoplasmic sample. The pellet was washed and then re-suspended in Nuclear Lysis Buffer (3mM EDTA, 0.2mM EGTA, 1mM DTT) and incubated on ice for 1h. Nuclear lysis was aided by sonication with a handheld homogeniser (VWR) for 10sec at 10min intervals. The nucleoplasmic supernatant and chromatin pellet were separated by centrifugation at 9000rpm for 10min at 4°C. The chromatin pellet was re-suspended in 200µl 2X Laemmli Buffer (Bio-Rad). Equal volumes of each fraction were used for Immunoblotting. Cytoplasmic and nucleoplasmic protein samples were diluted in 2X Laemmli Buffer and boiled for 5min at 95°C, then loaded on a 4-20% SDS-PAGE gel (Bio-rad) or a 3-8% Tris Acetate gel (Invitrogen). Proteins were wet transferred onto a PVDF membrane (Millipore) and assessed for successful transfer with Ponceau Red (Sigma). The membrane was blocked with 10% milk, 0.1% Tween-PBS for 20min, then incubated with primary antibodies in 1% milk, 0.1% Tween-PBS overnight at 4°C. Membranes were imaged with SuperSignal West Femto Maximum Sensitivity (Thermo) on an ImageQuant. Protein expression change or knock-down was quantified using ImageJ software to measure band intensity, normalised to loading control bands.

### Alkaline Phosphatase (AP) assay and quantification

Cells were seeded in 6-well plates and transfected with siRNAs at the time of plating as above. After 24hrs, cells were collected for RNA isolation and KD efficiency analysed by qRT-PCR. Cells from each condition were counted and 1,000 cells per well seeded into a new 6-well plate. Cells were re-transfected after 48hrs using 5pmol of siRNAs. Cells were fed every day. Four days after seeding cells at clonal density, the cells were assayed for alkaline phosphatase (AP) expression using StemTAG Alkaline Phosphatase staining kit (Cell Biolabs CBA-300). AP+ stained cells were imaged in 6-well plates using a M7000 Imaging System (Zeiss) with a 4X objective and a Trans-illumination brightfield light source. For quantification, area occupied by AP-high colonies was measured using ImageJ and plotted as fraction of total area of all colonies.

### RNA sequencing (RNA-seq) library preparation, sequencing, and analysis

Three biological replicates of mESCs were treated for 48hrs with control (Luciferase, siLuc) or Stag3 siRNAs, as above. Total RNA was isolated using NEB Monarch RNA prep kit (New England BioLabs). The RNA concentration and RIN (RNA integrity number) was assessed with a combination of NanoDrop 1000 (Thermo Scientific) and Agilent RNA 6000 Nano Bioanalyzer chip assay (Agilent). Libraries were prepared from 200ng total RNA using a KAPA mRNA HyperPrep kit (Kapa Biosystems, KK8580) and Roche SeqCap Adapter kit (Roche, KR1317) according to the manufacturer’s instructions. The libraries were quantified using a combination of Qubit dsDNA HS assay (Thermo Scientific) and Agilent DNA High Sensitivity Bioanalyzer chip assay (Agilent) and pooled together at equimolar concentrations. The final pooled library was quantified using a KAPA qRT-PCR Library Quantification assay (Kapa Biosystems, 07960140001). RNA-seq libraries were sequenced on a NextSeq 2000 platform (Illumina, Inc.) with a NextSeq 1000/2000 100 cycle P2 kit (Illumina, cat no. 20046811) using 66bp paired-end, single index reads.

Reads were quality controlled using FASTQC software. Transcript level count data (transcript length adjusted) was quantified using salmon (Patro et al., 2017) then imported and summarised in R at the gene-level into R using tximport (Soneson et al., 2015) and differential expression analysis performed using DESeq2 {Love:2014ka}. Gene Set Enrichment Analysis (GSEA) (Subramanian et al., 2005) was performed using the clusterProfiler package (Wu et al., 2021) using pathways from the Molecular Signature Database (MSigDB) (Castanza et al., 2023) accessed via the msigdbr package (https://cran.r-project.org/web/packages/msigdbr/vignettes/msigdbr-intro.html). Two custom gene sets were assayed in this study, ‘naïve pluripotency’, ‘primed pluripotency’ as previously described (Pezic et al., 2023). Gene sets were classed as having significant enrichment if the p-value was ≤0.05 and the normalised enrichment score (NES) exceeded +/- 1.

### Cell Cycle analysis and Annexin V staining

Cell cycle was assessed by flow cytometry of Hoechst staining. Confluent cells were resuspended in mES media at a concentration of 1x10^6^/ml supplemented with Hoechst 33342 (10μg/ml) (BD Bioscience, 561908) and incubated at 37°C for 45min. Cells were washed once in wash buffer (DMEM, 10% FCS), then re-suspended in FACS buffer for flow cytometry. Cell apoptosis was measured by Annexin V and Propidium Iodide (PI) staining. Confluent mES cells were washed twice in cell staining buffer (Biolegend 420201), then re-suspended in AnnexinV binding buffer at a concentration of 1x10^6^ cells/ml. 100µl aliquots were incubated with 5µl FITC Annexin V and 10µl PI solution (Biolegend 640914) for 15min at room temperature (25°C) in the dark. 400µl Annexin V Binding Buffer was added to each aliquot and staining profile was analysed by flow cytometry.

### Immunofluorescence and Microscopy

mESCs were cultured on fibronectin or gelatin-coated cover glass in 6-well plates. Cells were incubated in CSK buffer (10mM PIPES, 300mM sucrose, 100mM NaCl, 2mM MgCl2, 0.1% Triton X-100, 0.5mM PMSF, 1X Protease Inhibitor cocktail) for 4min at room temperature, then immediately fixed in 4% formaldehyde-PBS for 10min at room temperature before being washed and blocked in 10% FCS-PBS for 20min. Primary antibodies were diluted in 10% FCS, 0.1% Saponin (Sigma) and incubated overnight at 4°C. The next day, the cells were incubated with an Alexa Fluorophore-conjugated secondary antibody diluted in 10% FCS, 0.1% Saponin for 1hr at room temperature, washed and mounted on cover slides with ProLong Diamond Antifade Mountant with DAPI (Invitrogen P36962). Imaging of fixed cells was done using a LSM 880 confocal microscope (Zeiss) with a 63X oil objective. Analysis was performed using Fiji open-source software (ImageJ). Masks were applied using the ‘adjust threshold’ feature in Fiji in an unbiased manner to detect nuclei (DAPI) and centrosomes (gTub). IF intensity was quantified within the masked area.

### CRISPR-Mediated Stag3-v5 Knock-in Cell Line Generation

The v5 ssODN and guide RNA targeting *Stag3* 3’ terminal coding region was designed using Tagin Software (http://tagin.stembio.org) and purchased from IDT (sequences can be found in Table 2). Liophylysed gRNAs and ssODN were rehydrated in RNA duplex buffer (IDT 11-01-03-01) to 100μM and 30μM stocks, respectively. 2.2µl gRNA (100µM) was mixed with 2.2µl tracrRNA ATTO 550nm (100µM) (IDT) and annealed together using a controlled PCR step down cycle from 95°C-25°C (5°C every 20sec). The RNA duplex was then incubated with 20µg S.p Cas9 Nuclease V3 (IDT_1081058) for 10min at room temperature and stored on ice prior to transfection. 2.2µl v5 ssODN (30µM) sequence was mixed with 100% DMSO and denatured at 95°C for 5min. The ssODN was plunged immediately into ice. The RNP complex was mixed with 80,000 confluent 2i-grown mESCs re-suspended in P3 transfection buffer (Lonza V4XP-3032) before being transferred to an electroporation microcuvette well (Lonza V4XP-3032). Transfection was performed using a 4D Amaxa (Lonza) electroporator with the pre-programmed CG-104 setting. Post-nucleofection, the cells were seeded into a fibronectin-coated 6-well plate with fresh ES 2i media. The media was changed daily for four days without disturbing the cells, before being expanded into a T75 flask. Confluent ES cells were sparsely seeded into 10cm plates. Clones were manually picked into 96 well plates and expanded for selection by v5 IF, genotyping and Sanger sequencing.

### Chromatin Co-Immunoprecipitation (co-IP)

Cells were re-suspended in 0.1% NP-40-PBS (1ml/1x10^7^ cells) with 1X Protease Inhibitors (Roche) and 1mM DTT, and centrifuged at 1500rpm for 2min at 4°C. The pellet was re-suspended in Nuclear Lysis Buffer (3mM EDTA, 0.2mM EGTA, 1X Protease Inhibitors, 1mM DTT), vortexed for 30sec before being incubated on a rotator for 30min at 4°C and centrifuged at 6500g for 5min at 4°C to isolate the glassy chromatin pellet. This was re-suspended in High Salt Chromatin Solubilisation Buffer (50mM Tris-HCl pH 7.5, 1.5mM MgCl2, 300mM KCl, 20% glycerol, 1mM EDTA, 0.1% NP-40, 1mM Pefabloc, 1X Protease Inhibitors, 1mM DTT) with Benzonase (Sigma) (6U/1x10^7^) and incubated on rotator for 30min at 4°C. Chromatin was digested with 3x 10sec sonication at 30% intensity with a Vibra-Cell probe. The supernatant was collected by centrifugation at 1300rpm for 30min at 4°C, and then diluted to 200mM KCl concentration (IP buffer) with Dilution buffer (50mM Tris-HCl pH 7.5, 1.5mM MgCl_2_, 20% glycerol, 1mM EDTA, 0.1% NP-40, 1mM Pefabloc, 1X Protease Inhibitors, 1mM DTT). 30µl of A and G Dynabeads (Invitrogen) (15µl of each) were used per co-IP. Beads were washed 2x in 200mM KCl IP Buffer, re-suspended in IP Buffer with 10µg of the IP antibody, or an IgG-containing serum to match the species of the IP antibody and placed on rotator for 5h at 4°C. Beads were washed 3x in IP buffer and then incubated in 1mg chromatin lysate on a rotator overnight at 4°C. The beads were washed, re-suspended in 2X Laemmli Buffer (Bio-Rad), boiled for 10min at 95°C and used for WB as above.

### Whole Cell Extract Immunoprecipitation

An endogenous STAG3 IP from whole cell extract was initially attempted for Mass Spectroscopy. However, despite detection of known STAG3 interactors, no STAG3 peptides were identified. Briefly, 15µg anti-STAG3 or control rabbit IgG was conjugated to 30µl A and G Dynabeads beads (15µl each) as described above. WCE from 1x10^8^ mESCs was collected in 1ml RIPA buffer and 15mg of lysate was incubated with bead conjugates with rotation over night at 4°C. Flow-through was then removed, the beads were washed 5 times in RIPA buffer and eluted in 50µl formic acid (3%), which was incubated at 25°C for 5min with agitation. The resulting elute was prepared for Mass Spectrometry in the same way as described below for v5-TRAP IP/MS.

### v5-TRAP IP

CRISPR-targeted *Stag3*-v5 mESCs and WT mESCs were used for IP-mass spectrometry (IP-MS) where three v5-TRAP replicates were prepared per cell line. For each IP, 12.5x10^6^ confluent mESCswere harvested, washed once in ice-cold PBS-1% BSA and re-suspended in 500µl RIPA buffer (10mM Tris-HCl pH7.5, 150mM NaCl, 0.5mM EDTA, 2mM MgCl_2_, 1% NP40, 0.5% SDS, 0.5mM DTT, 1X EDTA-free Protease Inhibitors, 1mM Pefabloc, 1mM PMSF) with 50U Benzonase (Sigma). Lysate was vortexed for x3 10second pulses over 1min, then incubated on a rotator for 1h at 4°C. After 30min incubation, an additional 250U of heat-activated benzonase was added. Lysate was then centrifuged at 16000g for 10min. The supernatant was transferred to a 2ml tube containing 20µl magnetic Protein G-beads that had been previously washed in 1ml IP buffer (10mM Tris-HCl pH7.5, 150mM NaCl, 0.5mM EDTA, 2mM MgCl_2_, 0.05% NP40, 0.25% SDS). The remaining pellet was washed in 500µl Dilution Buffer and centrifuged again; this supernatant was added to the 2ml tube and the lysate was diluted with a further 1ml IP buffer (final SDS concentration 0.125%). The lysate was cleared of sticky proteins with Protein G-beads for 30min at 4°C on a rotator. A magnetic rack was used to extract the supernatant, which was transferred into a fresh 2ml tube containing 30 µl magnetic v5-TRAP beads (ChromoTEK) that had been previously washed in 1ml IP buffer. This was incubated for 1h10min at 4°C on a rotator. The supernatant was then removed as flow-through. The v5-TRAP beads were washed 3 times in 500µl wash buffer (10mM Tris-HCl pH7.5, 150mM NaCl, 0.5mM EDTA) and then 4 times with 500µl of 50mM triethylamonium bicarbonate (TEAB). On the final TEAB wash, the beads were transferred to a fresh 1.5ml tube. Proteins were eluted from the beads in 50µl formic acid (3%), which was incubated at 25°C for 5min with agitation. This elution was repeated once for a final IP elute volume of 100µl. The IP was vacuum-dried using a SpeedVac, washed once in 100µl TEAB (1M), then dried again before being re-suspended in 10µl TEAB (50mM) by vortexing and incubation at room temperature for 30min. 0.5µl (5mM) TCEP was added to the suspension and incubated at 37°C for 30min. 0.5µl (10mM) of Chloroacetamide was then added and the suspension was incubated in the dark for 20min at room temperature. Finally, the suspension was incubated at 37°C for 4h with 1.5µl (100ng/µl) trypsin. Resulting digested peptides were stored at -80°C until needed.

### Mass spectrometry and statistical data analysis

STAG3-v5 IP samples were analysed by liquid chromatography–tandem mass spectrometry (LC-MS/MS). nLC-MS/MS was performed on a Q Exactive Plus interfaced to a NANOSPRAY FLEX ion source and coupled to an Easy-nLC 1200 (Thermo Scientific). Samples were analysed as 30% and subsequently as 60% injections. Peptides were separated on a 27cm fused silica emitter, 75μm diameter, packed in-house with Reprosil-Pur 200 C18-AQ, 2.4μm resin (Dr. Maisch) using a linear gradient from 5% to 30% acetonitrile/ 0.1% formic acid over 30min, at a flow rate of 250nL/min. Peptides were ionised by electrospray ionisation using 1.8kV applied immediately prior to the analytical column via a microtee built into the nanospray source with the ion transfer tube heated to 320°C and the S-lens set to 60%. Precursor ions were measured in a data-dependent mode in the orbitrap analyser at a resolution of 70,000 and a target value of 3e6 ions. The ten most intense ions from each scan were isolated, fragmented in the HCD cell, and measured in the orbitrap at a resolution of 17,500.

Raw data were analysed with MaxQuant (Cox and Mann, 2008) version 1.6.17 where peptides were searched against the mouse UniProtKB database using default settings (http://www.uniprot.org/, downloaded 13/07/2018). Carbamidomethylation of cysteines was set as fixed modification, and oxidation of methionines and acetylation at protein N-termini were set as variable modifications. Enzyme specificity was set to trypsin with maximally two missed cleavages allowed. To ensure high confidence identifications, PSMs, peptides, and proteins were filtered at a less than 1% false discovery rate (FDR). Label-free quantification in MaxQuant was used with LFQ minimum ratio count set to 2 with ‘FastLFQ’ (LFQ minimum number of neighbours = 3, and LFQ average number of neighbours = 6) and ‘Skip normalisation’ selected. In Advanced identifications, ‘Second peptides’ was selected and the ‘match between runs’ feature was not selected. Statistical protein quantification analysis was done within the model-based statistical framework MSstats (Choi et al., 2014) (version 4.1) run through RStudio. Contaminants and reverse sequences were removed resulting in a STAG3-v5 interactome of 337 proteins and data was log2 transformed. To find differential abundant proteins across conditions, a linear mixed-effects model was fitted to the data.. The group comparison function was employed to test for differential abundance between conditions. Unadjusted p-values were used to rank the testing results and to define regulated proteins between groups. The group quantification function was used to obtain model-based log2 protein intensity summarisations across biological replicates. The data was plotted in a scatter plot shown in Figure 4a after filtering out the IgG peptide, uncharacterised proteins and proteins without values for log2 FC (this removed 10 proteins in total). The mass spectrometry proteomics data are deposited to the ProteomeXchange Consortium via the PRIDE (Perez-Riverol et al., 2022) partner repository with the dataset identifier PXD046456.

### Downstream Proteomic Analysis

Significantly depleted/enriched proteins were considered with an absolute log2foldchange >1 (2-fold change) and a p-value <0.05. STAG3-v5 interactome analysis was performed in STRING (https://string-db.org/). The network was generated as a full STRING network with a minimum interaction score of 0.7 and a high FDR stringency of 1%, resulting in 147 high confidence STAG3-v5 interactors, not including STAG3 itself. Over-enrichment of GO *Cellular Compartment*, *Biological Process* and *Molecular Function* terms were calculated with the mouse genome as background. Over-enrichment of protein motifs was calculated using the SMART domain database (http://smart.embl-heidelberg.de/), again, using mouse genome as background. The network shown in Fig 4 was manually rearranged in Cytoscape for visual clarity and displays all proteins from the high confidence STAG3-v5 interactors associated with the specified GO categories. Categories were visualized using the STRING pie chart function.

Custom proteome sets were generated from previously reported purified P-body and centrosome interactomes (Hubstenberger et al., 2017; O’Neill et al., 2022; Carden et al., 2023). Enrichment of our high confidence STAG3-v5 interactors in each set was analysed by hyperbolic distribution with significance calculated using dhyper function in R and multiple testing corrected for using the p.adjust Benjamini & Hochberg method, as previously described (Porter et al,. 2023). A total murine proteome of 54,822 proteins (https://www.uniprot.org/proteomes) was used to assess enrichment over background. Overlapping STAG3 and P-body/centrosome proteins were displayed using Cytoscope.

### TMT proteomics

Triplicate biological repeats of mESCs treated with *Stag3* and Luc siRNAs were prepared as described above. WCE was collected in RIPA buffer and quantified using Bradford Assay. 25µg of each sample was taken for TMT, and processed as described in (Azman et al., 2023), with some modifications. Briefly, samples were reduced by heating for 5min at 95°C in 50mM Bond-Breaker TCEP Solution (Thermo) and alkylated with 55mM of iodoacetimide for 30 minutes in the dark, before being diluted 8-fold in UA buffer (8 M urea, 100 mM Tris–HCl pH 8.8) and transferred to Vivacon 500 Hydrosart filters with a molecular cutoff of 30kDa. Samples were then concentrated by centrifugation at 14,000 g for 20min. The concentrated proteins were washed three times with 200μl of UA buffer through repeated buffer additions followed by centrifugations. This was followed by three washes with 200μl of 100 mM TEAB to remove the urea. Samples were then trypsin digested overnight at 37°C in a 600rpm shaking thermomixer, by adding 100μl of 100mM TEAB supplemented with 0.5μg of Trypsin (Sigma). Next day, TMT10plex label reagents were thawed, dissolved in acetonitrile, and added to each sample (8μg of TMT label per 1μg of input protein). The samples were incubated for 1h at 25°C, followed by quenching the unreacted TMT reagents with 5% hydroxylamine for 30min at 25°C. Peptides were then eluted by centrifugation at 14,000 g. Two additional elutions were then performed to maximise peptide extraction from the filter by adding 40μl of TEAB and centrifugation, plus a final elution with 40μl of 30% acetonitrile. Samples from each replicate set were then combined, along with a reference lysate channel, which was included in each TMT set. After combining all individually labeled eluates into one for each set, the pools were dried with a vacuum concentrator. The dried peptides were fractionated into seven fractions using Pierce™ High pH reverse-phase fractionation kit, according to the manufacturer’s instructions. The Fractions were dried again, before being reconstituted in A* buffer (0.1% TFA, 0.5% Acetic Acid, 2% Acetonitrile). LC– MS/MS analysis was performed on a Q Exactive-plus Orbitrap mass spectrometer coupled with a nanoflow ultimate 3000 RSL nano HPLC platform (Thermo Fisher), as described previously (Azman et al., 2023).

### Cyclohexamide chase experiments

Recovery of *Dppa3* mRNA in *Stag3* KD and control conditions was measured through inhibition of eEF2-mediated translation initiation with Cyclohexamide (CHX). Dppa3-T2A-GFP mESCswere treated with *Stag3* or Luc siRNAs as described above. At 48h, cell media was changed and supplemented with freshly prepared 100μg/ml CHX. Cells were incubated for 4h and 8h in CHX prior to harvesting. A no CHX-treated knockdown control was included in each experiment. GFP signal and cell viability in CHX-treated samples was measured by flow cytometry on a BD Fortessa X20 flow cytometer and analysed using FlowJo software (version 10.7.1). Total RNA was extracted from each condition as indicated above and *Dppa3* mRNA expression was assayed by qRT-PCR.

### Nascent translation analysis

For the Nascent translation analysis, Click-iT™ HPG Alexa Fluor™ 594 Protein Synthesis Assay Kit (Invitrogen C10429) was used. Cells were pre-incubated in Methionine-free media for 30min in the 37°C incubator before addition of L-homopropargylglycine (HPG) at 50µM. Cells were incubated with HPG for 30min, then collected, fixed, permeabilised, and stained using Click-It reaction in low retention tubes. HPG incorporation was measured by Flow Cytometry on a BD Fortessa X20. Analysis was performed using FlowJo software (version 10.7.1).

**Methods Table 1.** Antibodies used in this study.

| Protein | Catalogue No. | Company |
| --- | --- | --- |
| STAG3 (rabbit) | NBP1-58087 | Novus |
| STAG1 (goat) | ab4455 | Abcam |
| STAG2 (goat) | A300-158A | Bethyl |
| SMC3 (rabbit) | ab9263 | Abcam |
| RAD21 (rabbit) | ab992 | Abcam |
| NANOG (rabbit) | ab70482 | Abcam |
| alpha-TUBULIN (mouse) | T5168 | Sigma |
| gamma-TUBULIN (rabbit) | 15176-1-AP | Proteintech |
| gamma-TUBULIN (mouse) | ab27074 | Abcam |
| H3 (rabbit) | ab1791 | Abcam |
| V5 (mouse) | 14-6796-82 | Invitrogen |
| LAMIN B (goat) | Sc-6216 | Santa-Cruz |
| TNRC6C (mouse) | 101-11819 | RayBiotech |
| TNRC6C (rabbit) | orb186457 | Biorbyt |
| HORMAD1 (mouse) | 67091-1-Ig | Proteintech |
| V5-TRAP beads | v5tma | Proteintech |
| Donkey anti-mouse AF647 | A31570 | Invitrogen |
| Donkey anti-rabbit AF488 | A21206 | Invitrogen |
| Donkey anti-rabbit AF647 | A31573 | Invitrogen |
| Donkey anti-goat AF555 | A21432 | Invitrogen |
| Donkey anti-goat AF647 | A21447 | Invitrogen |
| Goat anti-Mouse AF568 | A-11031 | ThermoFisher Scientific |
| Goat anti-Rabbit AF647 | A-21244 | ThermoFisher Scientific |
| Rabbit anti-Goat AF647 | A-21446 | ThermoFisher Scientific |

**Methods Table 2.**
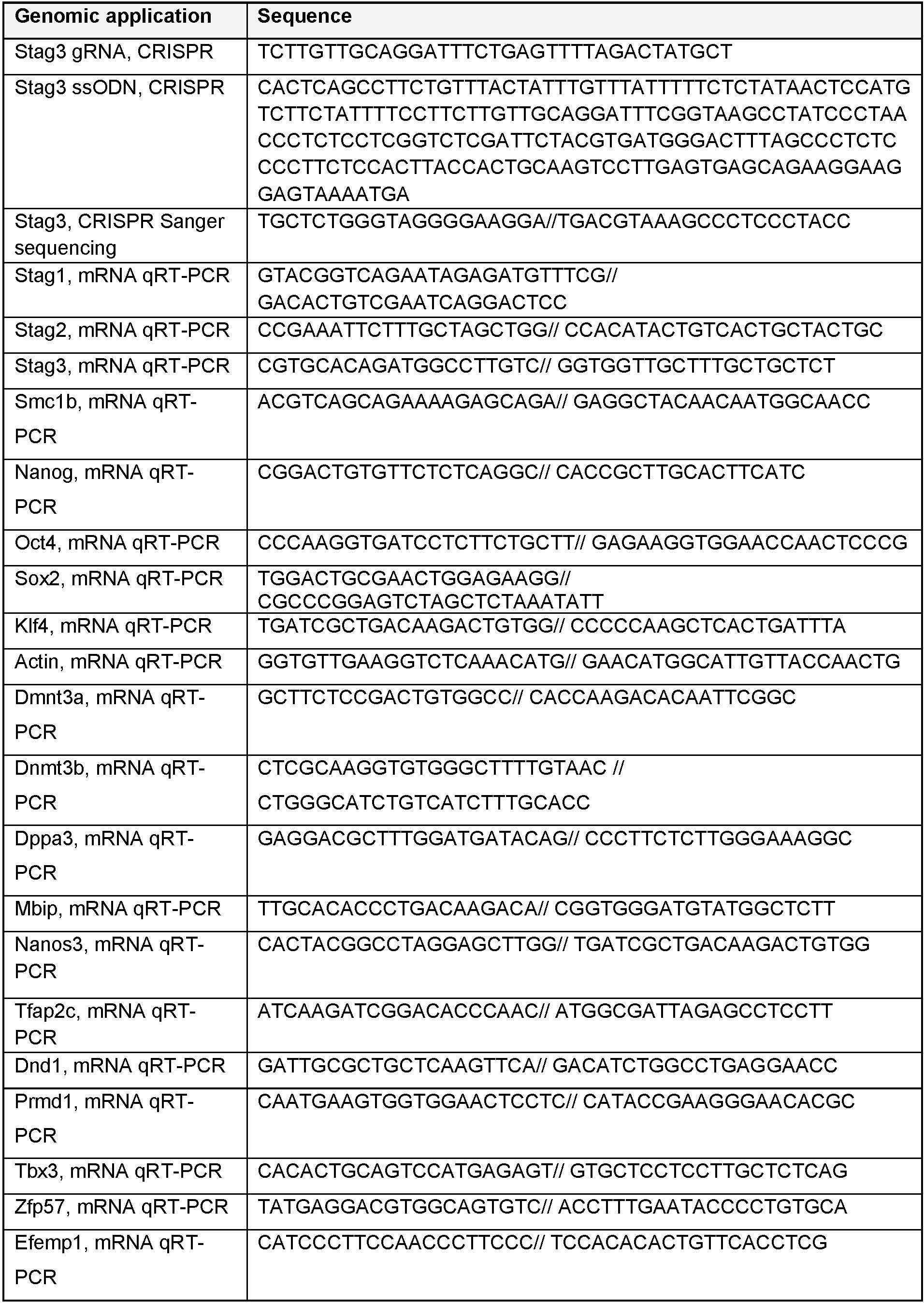
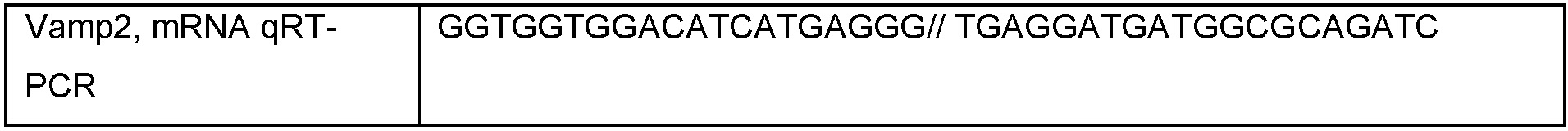
Oligos used in this study.

