## Supplementary information for "STAG3 promotes exit from pluripotency through post-transcriptional mRNA regulation in the cytoplasm"

### Supplementary Figure Legends

**Figure S1. Related to Figure 1 - mESCs require *Stag3* to exit pluripotency.**

a) Relative mRNA expression of the three *Stag* paralogs by qRT-PCR in serum-grown (FCS) mESCs. Data is from 10 independent biological replicates. NB. *Stag1* is ~5x more abundant than *Stag3*. The central line represents the median. Asterisks indicate a statistically significant difference as assessed using two-tailed t-test. *$p < 0.05$, **$p < 0.005$, ***$p < 0.0005$, ****$p < 0.0001$; ns, not significant.

b) WCE from mESCs analysed by IB for levels of STAG3 (as in Fig 1b) but without cropping the smaller bands (denoted by blue dots). SMC3, NANOG and alpha-TUBULIN (aTUB) are also shown.

c) Log2 fold change of *Stag* paralog mRNA levels, measured by qRT-PCR at several timepoints during EpiLC differentiation, relative to mESCs. *Nanog* and *Dnmt3a* are shown as down- and upregulated controls. Data is from 5 independent experiments.

d) Representative IB analysis of STAG3 levels in WCE of UT mESCs or after treatment with siLuc or siStag3. Blue dots denote bands that are sensitive to siStag3 and are likely isoforms. aTUB levels are shown as a loading control.

e) Left, representative FlowJo plots of the cell cycle based on FACS analysis of Hoechst-stained UT mESCs or after treatment with siLuc or siStag3. Right, Percentage of cells in each cell cycle stage in the three cell conditions, averaged from 3 independent experiments.

f) Representative FACS analysis of cell death based on Propidium iodide (PI) and AnnexinV staining of mESCs treated with siLuc or siStag3. *NB.* there was no significant difference in cell cycle distribution or cell death upon *Stag3* KD in mESCs compared to siLuc.

g) Relative expression of *Stag1, Stag2* and *Smc3* mRNA by qRT-PCR in UT mESCs or upon treatment with siLuc or siStag3. Data is from 9 independent biological replicates. Quantifications and statistical analysis as above.

h) WCE from mESCs analysed by IB for levels of STAG1 and STAG2 upon treatment with siLuc and siStag3. aTUB levels are shown as a loading control.

i) Relative expression of *Oct4, Sox2* and *Klf4* mRNA by qRT-PCR in UT mESCs or upon treatment with siLuc or siStag3. Data is from 9 independent biological replicates. Quantifications and statistical analysis as above.

j) WCE from mESCs analysed by IB for levels of NANOG upon treatment with siLuc and siStag3. H3 is shown as a loading control.

k) Relative expression of *Stag1, Stag2* and *Stag3* mRNA by qRT-PCR in mESCs treated with siLuc or siStag1. Data is from 5 independent biological replicates. Quantifications and statistical analysis as above. NB. *Stag2* and *Stag3* levels increase upon loss of the pro-pluripotency marker *Stag1*.

l) Area occupied by $AP^{hi}$ colonies relative to total colony area in mESCs treated with siLuc and siStag3 from three independent replicates where n>50 colonies/condition were counted and shown relative to UT cells.

**Figure S2. Related to Figure 2 - Loss of *Stag3* prevents commitment to PGCLC.**

a) Representative brightfield and epi-fluorescent images of PRDM1-GFP mESCs treated with siLuc or siStag3 at three timepoints of embryoid body (EB) differentiation. Scale bars represent 400um.

b) Representative FACS profile of PRDM1-GFP EBs at d2, 4 and 6 of differentiation either untreated (UT) or treated with siLuc or siStag3. Shown are the percentages of cells. Bottom, FlowJo analysis of siLuc (blue) and siStag3 (red) histograms.

c) Left, quantification of PRDM1-GFP GeoMean or right, the % of PRDM1-GFP+ cells assessed by FACS (relative to UT mESCs) at different stages of PGCLC differentiation and upon siRNA treatment. Data is from 5 independent biological replicates. Quantifications and statistical analysis as before.

**Figure S3. Related to Figure 3 - STAG3 is localised to the cytoplasm in mESCs.**

a) WCE from UT mESCs, or upon treatment with siLuc and siStag3, fractionated into cytoplasmic and chromatin fractions and analysed by IB for levels of STAG1, STAG2, STAG3 and RAD21. aTUB and H3 are shown as fractionation and loading controls.

b) Two independent chromatin immunoprecipitations of endogenous STAG3 in WT mESCs using a commercial STAG3 antibody and Mock control. Input (INP) is 2.5% of lysate. Top, STAG3 interacts with SMC3 and bottom, RAD21 and CTCF on chromatin. Arrows indicate likely STAG3 isoforms as indicated in Figure 3.

c) Immunofluorescence (IF) controls in mESCs and representative STAG3 staining in siLuc and siStag3 treated cells.

d) coImmunofluorescence (coIF) of the three STAG paralogs in mESCs alongside F-actin. *NB* the primarily nuclear localisation of STAG1 and STAG2 but relative absence of nuclear signal for STAG3.

e) coIF of STAG1, STAG3 and RAD21 in mESCs. *NB* RAD21 signal appears to be absent from STAG3 signal at distinct cytoplasmic puncta and at the spindle.

f) Top, Schematic of CRISPR *knock-in of a v5 sequence at the 3*' region of *Stag3.* Bottom, PCR amplification of *Stag3* 3' region from clones used in this study. *NB* clone C3's doublet band represents heterozygote v5 knock-in.

g) Immunoblot of cytoplasmic, nucleoplasmic and chromatin fractions from UT, siLuc or siStag3 treated *Stag3-v5* mESCs. Blue and red dots indicate canonical and band-shifted STAG3 or STAG3-v5, respectively. aTUB and H3 are shown as fractionation and loading controls.

h) Top, coIF of STAG3-v5, LAMIN B and gTUB. STAG3 is detected here using a v5 antibody. Scale bars, 3um. Bottom, quantification of IF results. Data are from one biological replicate where > 100 independent cells/condition. See also Figure 3.

**Figure S4. Related to Figure 4 - Characterization of the STAG3 protein interaction network in mESCs.**

a) Scatter plot displaying the Log2 protein intensity of all the 337 proteins identified in STAG3-v5 IP-MS data produced from three biological replicates of v5-TRAP in WT and *Stag3*-v5 mESCs. Labelled are the ribosome proteins that were not included in the main Figure 4a for clarity. Red dots represent statistically significantly enriched STAG3-v5 interactors with changes of at least 2-fold and $p<0.05$.

**Figure S5. Related to Figure 5 - STAG3 mediates post-transcriptional regulation of *Dppa3* in mESCs and destabilises TNRC6C.**

a) Analysis of global levels of nascent translation by measuring HPG incorporation using Flow cytometry and analysed using FlowJo software. Shown is the quantification of the change in HPG incorporation relative to UT mESCs in both the high and low HPG population. Data are from 4 independent replicates.

b) Relative qRT-PCR mRNA expression of select genes in siLuc and siStag3 mESCs treated with cyclohexamide (CHX) for 4 or 8 hours to inhibit translation from. Data is shown as an average from two independent replicates and relative to siLuc, no CHX samples. Genes were selected based on their mRNA down-regulation (RNA-seq, Fig 1e) and protein up-regulation (TMT, Fig 5b).

c) Further representative coIFs of STAG3, LAMINB and TNRC6C in WT ESCs. See also Figure 5.

**Supplementary Figure 1** *Weeks et al.*

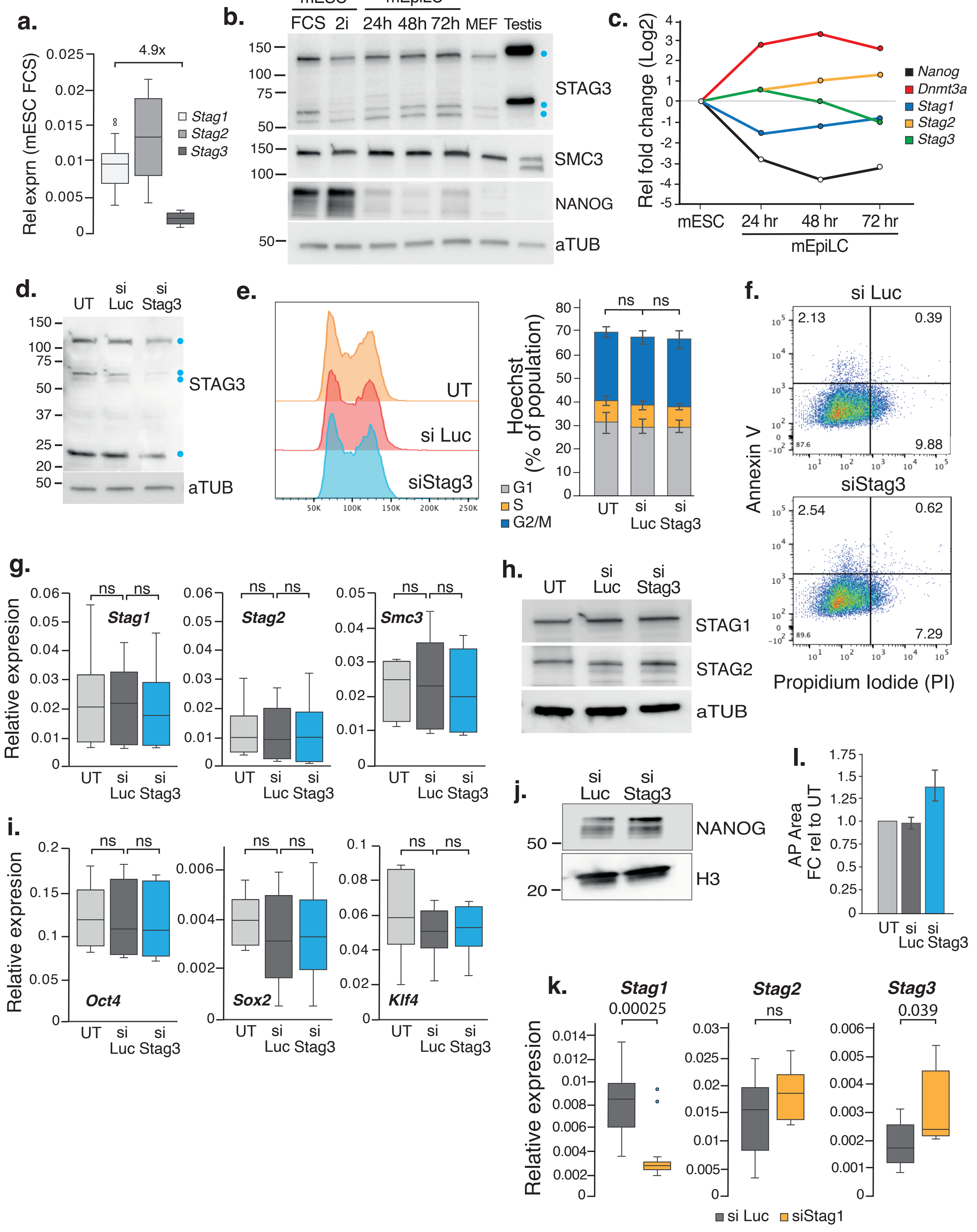

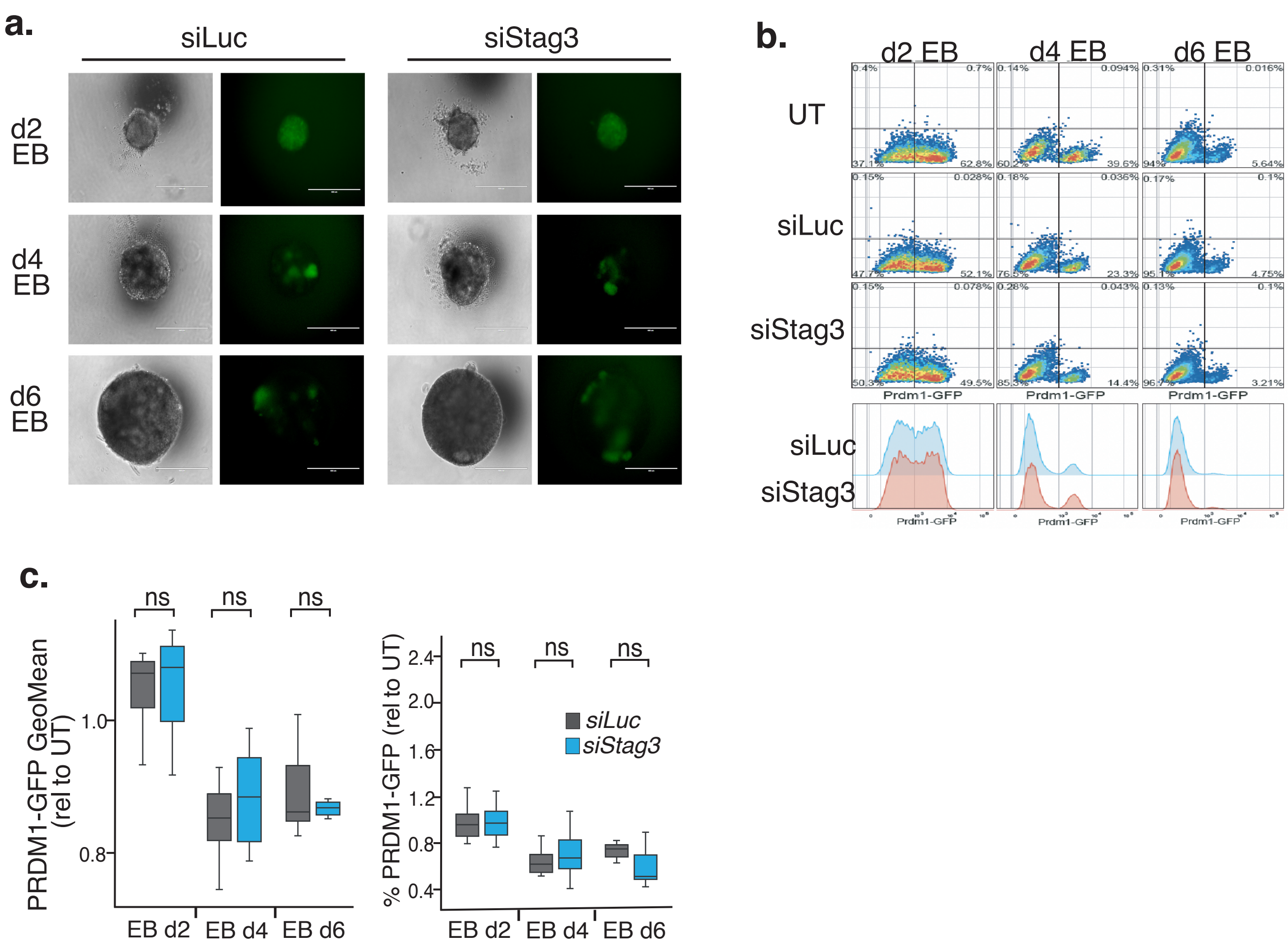
a.
siLuc
siStag3
d2 EB
d4 EB
d6 EB
b.
d2 EB
d4 EB
d6 EB
UT
siLuc
siStag3
Prdm1-GFP
siLuc
siStag3
c.
ns
PRDM1-GFP GeoMean (rel to UT)
% PRDM1-GFP (rel to UT)
EB d2
EB d4
EB d6
siLuc
siStag3

### Supplementary Figure 3

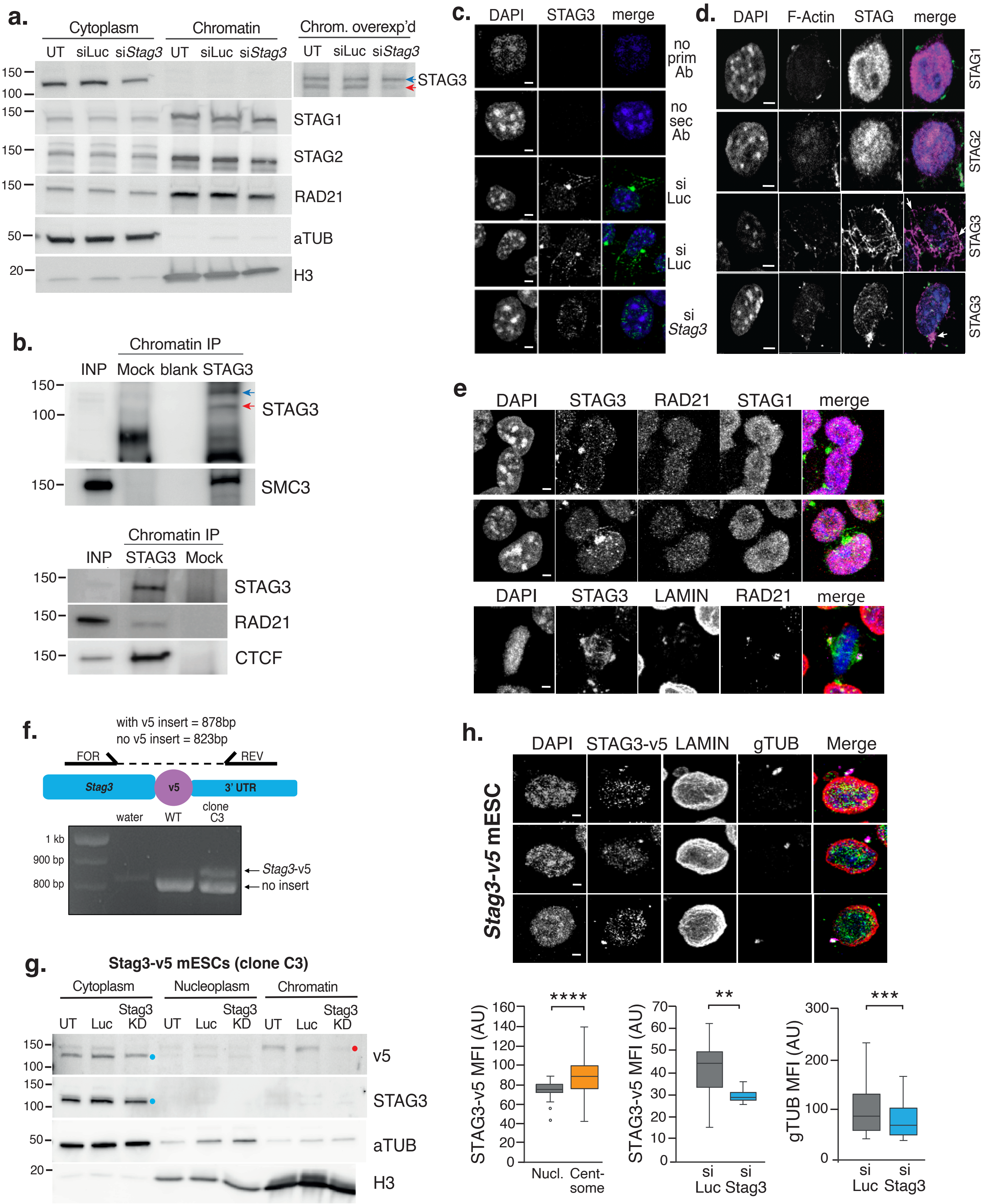

**a.**

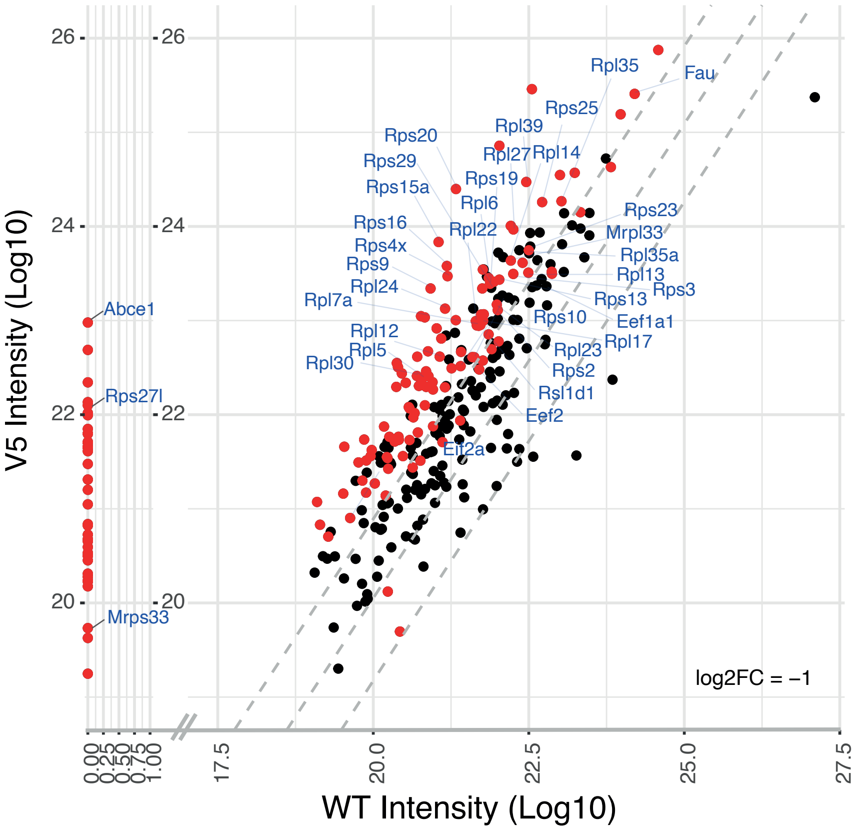
V5 Intensity (Log10)
WT Intensity (Log10)
log2FC = −1
Rpl35
Fau
Rps25
Rps20
Rpl39
Rps29
Rpl27
Rpl14
Rps15a
Rps19
Rpl6
Rps23
Rps16
Rpl22
Mrpl33
Rps4x
Rpl35a
Rps9
Rpl13
Rpl24
Rps3
Rpl7a
Rps13
Rps10
Eef1a1
Rpl12
Rpl17
Rpl5
Rpl23
Rpl30
Rps2
Rsl1d1
Eef2
Eif2a
Abce1
Rps27l
Mrps33
26
24
22
20
0.00
0.25
0.50
0.75
1.00
17.5
20.0
22.5
25.0
27.5

### Supplementary Figure 5

*Weeks et al.*

**a.**

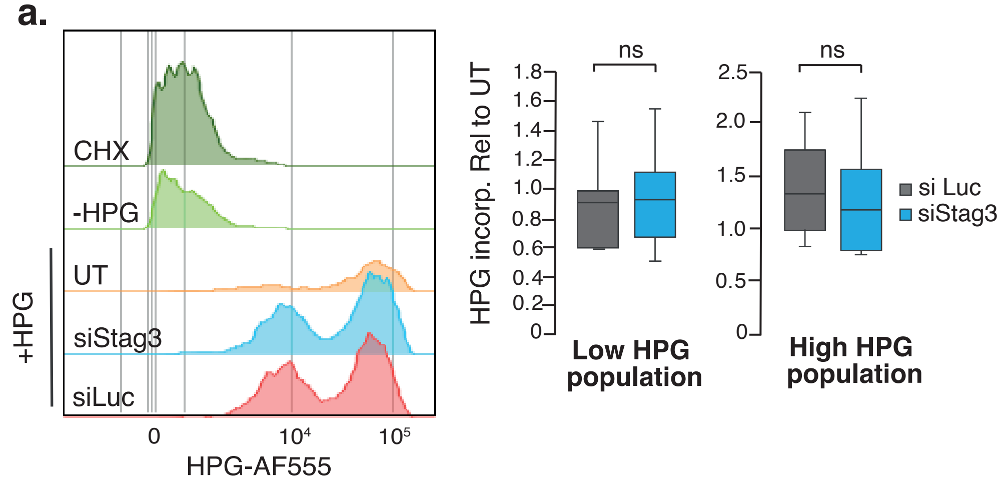


**b.**

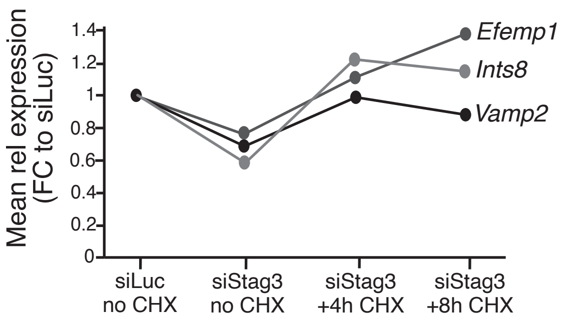


**c.**

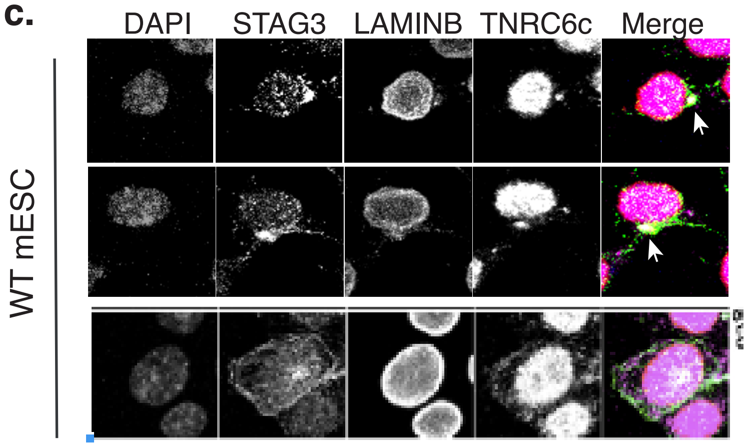
